# RhoB promotes *Salmonella* survival by regulating autophagy

**DOI:** 10.1101/2023.04.05.535690

**Authors:** Marco Kirchenwitz, Jessica Halfen, Kristin von Peinen, Silvia Prettin, Jana Kollasser, Cord Brakebusch, Klemens Rottner, Anika Steffen, Theresia E.B. Stradal

## Abstract

*Salmonella enterica* serovar Typhimurium manipulates cellular Rho GTPases for host cell invasion by effector protein translocation via the Type III Secretion System (T3SS). The two Guanine nucleotide exchange (GEF) mimicking factors SopE and –E2 and the inositol phosphate phosphatase (PiPase) SopB activate the Rho GTPases Rac1, Cdc42 and RhoA, thereby mediating bacterial invasion. *S.* Typhimurium lacking these three effector proteins are largely invasion-defective. Type III secretion is crucial for both early and later phases of the intracellular life of *S.* Typhimurium. Here we investigated whether and how the small GTPase RhoB, known to localize on endomembrane vesicles and at the invasion site of *S.* Typhimurium, contributes to bacterial invasion and to subsequent steps relevant for *S.* Typhimurium lifestyle.

We show that RhoB is significantly upregulated within hours of *Salmonella* infection. This effect depends on the presence of the bacterial effector SopB, but does not require its phosphatase activity. Our data reveal that SopB and RhoB bind to each other, and that RhoB localizes on early phagosomes of intracellular *S.* Typhimurium. Whereas both SopB and RhoB promote intracellular survival of *Salmonella*, RhoB is specifically required for *Salmonella*-induced upregulation of autophagy. Finally, in the absence of RhoB, vacuolar escape and cytosolic hyper-replication of *S.* Typhimurium is diminished. Our findings thus uncover a role for RhoB in *Salmonella*-induced autophagy, which supports intracellular survival of the bacterium and is promoted through a positive feedback loop by the *Salmonella* effector SopB.

## Highlights

- The *Salmonella* effector SopB induces expression of host cell RhoB
- RhoB expression and SopB secretion enhance each other in a positive feedback loop
- RhoB deficiency results in poor survival of *Salmonella*
- *Salmonella*-induced autophagy is abrogated in cells lacking RhoB

## Introduction

*Salmonella enterica* serovar Typhimurium is a Gram-negative, food borne pathogen causing gastroenteritis in humans. This pathogen has developed a diverse set of virulence factors that trigger host cell signaling leading to cytoskeletal rearrangements and inflammatory responses, subsequently promoting infection, bacterial replication and subcellular distribution (Finlay & Brumell, 2000; Haraga, Ohlson, & Miller, 2008). *S.* Typhimurium expresses a macromolecular complex present on its surface termed type III secretion system (T3SS), embodying a needle-like complex facilitating the translocation of bacterial effector proteins into the host cell cytoplasm. T3SSs are a common and conserved virulence strategy of many Gram-negative pathogens. Whereas the cocktail of translocated effectors can differ among distinct pathogens, they commonly harbor factors that hijack the host’s Rho GTPase signaling network. This enables them to manipulate the host’s actin cytoskeleton for bacterial adhesion, invasion and to eventually establish their own niche (Alto et al., 2006; Stradal & Schelhaas, 2018). Despite significant progress on this topic in recent years, such host-pathogen interfaces are complex and still not fully understood.

Major drivers triggering *S.* Typhimurium invasion into non-phagocytic cells are the effector proteins SopB and SopE translocated by the *Salmonella* pathogenicity island-1 (SPI-1) encoded T3SS (T3SS-1) (Kubori et al., 1998; Mills, Bajaj, & Lee, 1995). More specifically, the effectors SopE and its paralogue SopE2 are bacterial guanine exchange factors (GEFs) that activate the host Rho GTPases Rac1, Cdc42 and RhoG (Friebel et al., 2001; Hanisch et al., 2011; Hardt, Chen, Schuebel, Bustelo, & Galan, 1998; Stender et al., 2000), and induce membrane ruffling followed by bacterial uptake on the pinocytic route. In addition, SopB mediates RhoA/myosin II-dependent invasion by inducing contractility (Hanisch et al., 2011). SopB harbors a phosphoinositide phosphatase (PIPase) activity hydrolyzing PI(3,4)P_2_ and PI(3,4,5)P_3_ inside the host cell, indirectly inducing Cdc42 and RhoG activation (Norris, Wilson, Wallis, Galyov, & Majerus, 1998; Patel & Galan, 2006) and activation of AKT signaling. This in turn promotes intracellular survival by inhibiting cellular apoptosis (Knodler, Finlay, & Steele-Mortimer, 2005). Finally, the N-terminus of SopB was shown to bind to Cdc42 and mutation of this region impaired intracellular replication (Rodriguez-Escudero, Ferrer, Rotger, Cid, & Molina, 2011). After cell invasion, *S.* Typhimurium remains in the phagosome and secrets effector proteins via a second T3SS (T3SS-2) encoded on a second pathogenicity island, SPI-2. This leads to rearrangements of the endomembrane system and matures the phagosome into the *Salmonella* containing vacuole (SCV) (Drecktrah, Knodler, Howe, & Steele-Mortimer, 2007; Hensel et al., 1998; Jennings, Thurston, & Holden, 2017), the intracellular replication niche of *S.* Typhimurium (Beuzon et al., 2000). Some bacteria escape the SCV and reach the host cytosol where they can also replicate, which depends on the T3SS-1 but not T3SS-2 (Knodler, Nair, & Steele-Mortimer, 2014; Knodler et al., 2010; Malik-Kale, Winfree, & Steele-Mortimer, 2012). During *Salmonella* invasion, the host cell GTPase RhoB is recruited to sites of infection, which was described to depend on the effector protein SopB (Truong et al., 2018), although the function of this recruitment has remained elusive. Recently, RhoB was reported to play a role in the regulation of autophagy in normal and infected cells (M. Liu et al., 2018; Miao et al., 2021). RhoB can be posttranslationally modified through palmitoylation and prenylation at Cys189 and 193, respectively, which determines its subcellular localization (Adamson, Marshall, Hall, & Tilbrook, 1992; Michaelson et al., 2001; Wherlock, Gampel, Futter, & Mellor, 2004). Moreover, RhoB localizes at the plasma membrane and in addition, unlike RhoA and –C, at endosomal and pre-lysosomal vesicles (Michaelson et al., 2001), regulating transport processes (Fernandez-Borja, Janssen, Verwoerd, Hordijk, & Neefjes, 2005).

Here, we describe how RhoB contributes to *S.* Typhimurium invasion and intracellular survival. RhoB was reported to be recruited to *S.* Typhimurium entry sites at the host cell plasma membrane. However, a role during later stages of infection when *S.* Typhimurium interacts with endomembranes, where RhoB is prominently localized in cells, has not been described. In this study, we show that RhoB is upregulated in expression in the first hours after *Salmonella* invasion, which depended on the bacterial effector SopB. However, in contrast to SopB’s role in RhoA and contractility activation during invasion, SopB-mediated upregulation of RhoB does not require the phosphatase activity of SopB. Moreover, our data reveal that RhoB not only localizes on the SCVs of intracellular *S.* Typhimurium, but also that SopB and RhoB bind to each other. Whereas both RhoB and SopB promote intracellular survival of *Salmonella*, only RhoB but not SopB is required for *Salmonella*-induced upregulation of autophagy. Our findings point towards a role of RhoB in *Salmonella*-induced autophagy and directly or indirectly promoting *Salmonella* survival.

## Materials and Methods

### Cultivation of cells and transfections

*Shigella flexneri* M90T 5a (Sansonetti, Kopecko, & Formal, 1982), *Salmonella enterica* serovar Typhimurium strain SL1344 WT (Hoiseth & Stocker, 1981) and its isogenic SopB-deleted strain (ΔSopB) (Hanisch et al., 2011) bearing spectinomycin resistance were cultured in Luria-Bertani (LB) broth at 37 °C. ΔSopB was grown in the presence of 100 µg/ml spectinomycin, and *Salmonella* expressing TEM-tagged effectors were grown in the presence of 50 µg/ml kanamycin. *E. coli* transformed with the respective plasmids for plasmid amplification (Supplemental Table 1) were grown in the presence of appropriate antibiotics (ampicillin, 100 µg/ml; chloramphenicol, 20 µg/ml; kanamycin, 50 µg/ml; tetracycline, 15 µg/ml; and spectinomycin, 100 µg/ml).

NIH/3T3 fibroblast cells (ATCC CRL-1658) and respective RhoB knockout clones (generated in this study) were cultured in DMEM with 4.5 g/l glucose, 10 % fetal bovine serum, 1 % glutamine and 1 % MEM-non-essential-amino-acids at 37 °C and 7.5 % CO_2_. Cells were transfected with X-tremeGENE 9 DNA Transfection Reagent (Roche, Basel, Switzerland) following manufacturer’s instructions.

### Plasmids and Cloning

Bacterial expression plasmids were generated via Gateway cloning (Invitrogen, Carlsbad, CA, USA). The effector sequences of SopB, SopB^C460S^ and SopE1 from *S.* Typhimurium, IpgB2 and IpgD from *Shigella flexneri* were custom synthesized by Genscript (Piscataway, NJ, USA) and for further cloning amplified by nested PCR using gateways cloning primers. All primers are listed in Supplemental Table 1. These gene fragments were cloned into pDONR223 (Invitrogen) by gateway cloning as described previously (Hartley, Temple, & Brasch, 2000; Rual et al., 2004). For translocation assays, the IPTG-inducible vector pK184-ccdB-TEM1 was generated. First, the β-lactamase gene TEM-1 was amplified from plasmid pCX340 (Charpentier & Oswald, 2004) and cloned into pK184 plasmid (Jobling & Holmes, 1990) using BamHI and HindIII restriction enzyme sites. Subsequently, the gateway cassette was cloned into pK184-TEM1 vector using the SmaI restriction enzyme. Finally, SopB and SopE were cloned from pDONR233 vectors into pK184-ccdB-TEM1 via gateway cloning, as previously described (Katzen, 2007). For effector expression in mammalian cells, the gateway cassette of the plasmid pmCherry-ccdB was generated by subcloning of ccdB into pmCherry-C1 plasmid (Clontech, Mountain View, CA) using SmaI. The effectors SopB, SopB^C460S^, SopE1, IpgB2 and IpgD were cloned into pmCherry-ccdB via gateway cloning. For pRK5-myc-SopB generation, SopB was amplified from pDONR223-SopB using SopB_pRK5_fw and SopB_pRK5_rev primers, and cloned into pRK5-myc plasmid using BamHI and HindIII restriction enzymes. Human pRK5-myc-IRSp53 and pFS48 (expressing mCherry) plasmid were as described (Disanza et al., 2006; Nuss et al., 2016). Plasmids of this study are listed in Supplementary data 1. All generated plasmids were sequence-verified.

### Generation of *S.* Typhimurium SopB^C460S^ mutant by homologous recombination using the λ–Red system

The SopB^C460S^ mutation was introduced in *S.* Typhimurium genome by allelic replacement using the λ-Red system as previously described (Datsenko & Wanner, 2000). Briefly, a tetracycline resistance gene cassette with 40 bp of flanking homologous regions of SopB was amplified by PCR using tetRA_SopB_fw and tetRA_SopB_rev primers. This gene fragment was transformed into *S.* Typhimurium wild-type expressing the λ-Red recombinase on pKD46 (Datsenko & Wanner, 2000). Subsequently, bacteria were counter-selected on LB-medium with 15 µg/ml tetracycline (TetR) and tetracycline-sensitive medium (TetS) (5 g/l tryptone, 5 g/l yeast extract, 12 g/l agar, 10 g/l NaCl, 10 g/l NaH_2_PO_4_ x H_2_O, 0.1 mM ZnCl_2_, 12 mg/l fusaric acid, 1 mg/l anhydrotetracycline) for tetracycline-resistant mutants. For allelic replacement of the tetracycline resistance gene cassette, the SopB^C460S^ gene fragment was inserted in fusaric acid-sensitive and tetracycline-resistant *S.* Typhimurium expressing λ-Red recombinase on pKD46. Bacteria were counter-selected on TetR and TetS medium and selected for loss of tetracycline resistance. All clones were verified by PCR and sequencing.

### Generation of RhoB KO cell lines with CRISPR/Cas9

RhoB KO cells were generated by using the CRISPR/Cas9 technology. pSpCas9(BB)-2A-GFP (PX458) was from Feng Zhang (Addgene plasmid # 48138; http://n2t.net/addgene:48138; RRID:Addgene_48138) (Ran et al., 2013). The single guide RNA (sgRNA) sequence targeting RhoB was 5’-GCACCACCAGCUUCUUGCGGA-3’, and cloned into pSpCas9(BB)-2A-GFP. NIH/3T3 cells were transfected with pSpCas9(BB)-2A-GFP:sgRNA-RhoB and single GFP-positive cells sorted into 96-well plates via FACS using Aria-II SORP (BD Biosciences, Heidelberg, Germany). Cells were expanded and single clones screened for the absence of RhoB expression via immunoblotting. Genotypes of potential KO clones were determined via sequencing, as previously described (Kage et al., 2017). RhoB was amplified using RhoB_clon_fw and RhoB_clon_rev primers. Amplified fragments were subcloned into pCR4Blunt-TOPO vector (Zero Blunt TOPO PCR Cloning Kit, Invitrogen) and sequenced with primers M13_seq_fw and M13_seq_rev. Sequences were analyzed for frameshift mutations and monoallelic or biallelic deletions and insertions. All three single cell-derived clones were verified for loss of RhoB expression by Western blotting. RhoB KO cells used in experiments represented a pool of three independently grown clones pooled prior to each experiment.

### Gentamycin protection assays

For analysis of *S.* Typhimurium invasion and survival, gentamycin protection assays were performed, as previously described (Kirchenwitz et al., 2022). Briefly, 5×10^4^ cells per well were seeded into 24-well plates and incubated at 37 °C in a humidified, 7.5 % CO_2_ atmosphere for 24 h. Fresh LB broth was inoculated with overnight culture of *S.* Typhimurium and grown at 37 °C under agitation up to an OD_600_ 0.8 ∼ 1.0. Subsequently, bacterial suspension was harvested at 3000 x g for 2 minutes and diluted in DMEM to a MOI of 100. Infection was initiated by centrifugation of plates after addition of *S.* Typhimurium for 5 min at 935 x g. After 30 min incubation time, 50 µg/ml gentamycin in DMEM was added. Cells were infected for different time points as indicated in Figures, washed thrice with PBS, lysed with 0.5 % Triton X-100 for 5 min on ice and diluted in PBS for plating onto agar plates. Plates were incubated overnight at 37 °C and then colonies were counted using an automated colony counter (Scan4000, Interscience, Saint Nom la Brétèche, France).

### Adhesion Assay

For adhesion assays, cells and bacteria were prepared as described before (see gentamycin protection assays). After centrifugation of bacteria onto host cells, dishes were incubated for 15 min, washed thrice with pre-warmed DMEM and either directly lysed, or incubated in 50 µg/ml gentamycin in DMEM for 45 min and then lysed and processed as described above. The rate of adhesion was determined by subtracting the number of intracellular bacteria (1 hpi) from adhesive bacteria (15 min pi).

### Real-time effector translocation assay

Overnight cultures of *S.* Typhimurium expressing pK184-SopB-TEM1 and pk184-SopE-TEM1, respectively, were grown in LB-medium supplemented with kanamycin and 1 mM IPTG. NIH/3T3 cells were seeded in a 96-well plate at a density of 30.000 cells/well. Fresh LB-medium supplemented with 1 mM IPTG, 0.3 M NaCl and kanamycin was inoculated with overnight bacterial cultures and grown for 3 h until OD_600_ between 0.8 and 1.0 was reached. Cells were incubated with DMEM containing CCF4-AM loading solution (Thermo Fisher Scientific, K1029, Waltham, MA, USA) and 2.5 mM probenecid 1.5 h prior to infection. Bacteria were collected by centrifugation at 3000 x g for 3 min and adjusted to an MOI of 300 containing 1 mM IPTG and 2.5 mM probenecid. Infection was initiated by addition of *S.* Typhimurium to cells and centrifugation for 5 min at 935 x g. Cells were immediately transferred and imaged by spinning disk microscopy at 37 °C in a humidified 7.5 % CO_2_ atmosphere as described below. Cells were imaged using 405 nm excitation combined with 447/60 and 525/50 nm emission filters at 10 min intervals. After 30 min of infection, 50 µg/ml gentamycin in DMEM was added. Fluorescence intensities of different time points were analyzed with NIS-Elements (Nikon, Düsseldorf, Germany). The rate of effector translocation was determined by calculating the ratio of the fluorescence intensity of the product (447 nm) and the fluorescence intensity of the substrate (525 nm).

### Cell lysates, protein measurements and Western Blotting

Protein samples were prepared by washing cells thrice with ice-cold PBS and lysing with ice-cold lysis buffer (50 mM Tris HCl, pH 8, 10 % glycerol, 100 mM NaCl, 20 mM NaF, 4 mM Na_3_VO_4_, 1 % Nonidet P-40, 2 mM MgCl_2_, 1x protease inhibitor (A32955, Thermo Fischer Scientific) or 4x SDS-Laemmli buffer as indicated in figure legends. Analysis of proteins by SDS-PAGE and immunoblotting was performed according to standard procedures. Primary antibodies used are GAPDH (clone 6C5, #CB1001, 1:5000 dilution, RRID: AB_1285808, Merck Millipore, Burlington, MA, USA), GFP (ab290, 1:2000 dilution, RRID: AB_303395, Abcam, Cambridge, UK), Myc-Tag (2276, 1:1000 dilution, RRID: AB_331783, Cell Signaling Technology, Danvers, MA, USA), RhoA (sc-418, Clone 26C4, 1:200 dilution, RRID: AB_628218, Santa Cruz, Dallas, TX, USA), RhoB (sc-180, Clone 119, 1:1000 dilution, RRID: AB_2179110, Santa Cruz), RhoB (sc-8048, Clone 5, 1:1000 dilution, RRID: AB_628219, Santa Cruz), RhoC (3430, 1:1000 dilution, RRID: AB_2179246, Cell Signaling Technology, Danvers, MA, USA), caspase-3 (9662, 1:1000 dilution, RRID: AB_331439, Cell Signaling Technology), cleaved-caspase-3 (9661, 1:1000 dilution, RRID: AB_2341188, Cell Signaling Technology), AKT (9272, 1:1000 dilution, RRID: AB_329827, Cell Signaling Technology), phospho-AKT (Ser473) (9271, 1:1000 dilution, RRID: AB_329825, Cell Signaling Technology), LC3A/B (4108, 1:2000 dilution, AB_2137703, Cell Signaling Technology,) and SQSTM1/p62 (ab56416, 1:2000 dilution, RRID: AB_945626, Abcam). Primary antibodies were detected by peroxidase-conjugated goat α-mouse and goat α-rabbit antibodies (115-035-062 and 111-035-045, 1:5000 dilution, RRID: AB_2338504 and AB_2338504, Jackson ImmunoResearch, Cambridge, UK) using the Lumi-Light Western Blotting substrate (Roche) with the ECL Chemocam IMAGER (Intas, Göttingen, Germany). Proteins were analyzed by densitometric analysis and normalized to the levels of GAPDH.

### RT-qPCR

For mRNA expression analysis, cells were lysed and mRNA was extracted using NucleoSpin RNA Plus kit (Macherey-Nagel, Dueren, Germany) kit and stored at −80°C. Samples were analyzed by real-time, quantitative PCR (RT-qPCR) with the SensiFAST SYBR No-ROX One-Step Kit in a LightCycler 96 (Roche) according to the manufacturer’s instructions. The primers for murine RhoB were ordered from Sino Biological (MP202540, China). The sequence of primers for RPS9 were 5’-CTGGACGAGGGCAAGATGAAGC-3’ and 5’-TGACGTTGGCGGATGAGCACA-3’ (Eurofins Scientific, Luxembourg). Reactions were incubated at 45 °C for 10 min and 95 °C for 2 min, followed by 45 cycles of 95 °C for 5 sec, 60 °C for 10 sec and 72 °C for 5 sec. Changes in gene expression were calculated using the 2^−ΔΔCT^ method.

### Immunoprecipitation

Cells were washed with ice-cold PBS, lysed in lysis buffer, and subjected to sonification. Cleared lysates were incubated with anti-GFP antibody overnight at 4 °C with constant rotation, and subsequently incubated with equilibrated magnetic beads (Dynabeads Protein G, Thermo Fisher Scientific) for 2 h at 4 °C with constant rotation. Beads were separated with magnets and washed thrice with lysis buffer. Proteins were eluted with 4x SDS-Laemmli buffer, boiled and analyzed by immunoblotting.

### Annexin V Apoptosis assay

NIH/3T3 wild-type and RhoB KO cells were seeded into a 96-well plate (flat bottom) at a density of 17.000 cells/well and incubated overnight. Cells were washed twice with pre-warmed DMEM, incubated for 30 min and infected with *S.* Typhimurium wild-type and ΔSopB at MOI 100 for 30 min as described above. Infection was synchronized by centrifugation at 935 x g. Subsequently, cells were washed twice with DMEM and incubated in growth medium containing 2.5 mM CaCl_2_ and Alexa Fluor 488-conjugated annexin V (Thermo Fisher Scientific). Cells were imaged and analyzed in an Incucyte S3 live-cell analysis system (Sartorius, Göttingen, Germany), using the adherent cell-by-cell module to assess phase contrast and green and red fluorescence intensities.

### Immunofluorescence

For immunofluorescence analysis, cells were seeded onto 25 µg/ml fibronectin-coated coverslips, as previously described (Steffen, Kage, & Rottner, 2018). Cells were fixed with pre-warmed 4% PFA (paraformaldehyde) in PBS for 20 min at room temperature and incubated with 0.5% Triton X-100 in PBS for 5 min. After blocking with 5 % horse serum in 1 % BSA in PBS for 30 min, cells were stained with LC3 (M152-3, 1:200 dilution, RRID: AB_1279144, MBL International, Woburn, MA, USA) antibodies for 16 h at 4°C, washed with PBS and labeled with secondary antibody for 1 h. Staining of F-actin was performed with phalloidin-Alexa Fluor-488 (A12379, 1:300 dilution, RRID: AB_2759222, Thermo Fischer Scientific). Cells were mounted on glass slides with ProLong Diamond (P36971, Thermo Fisher Scientific).

### LysoTracker staining

For SCV staining, NIH/3T3 wild-type and RhoB KO cells were seeded onto fibronectin-coated coverslips. Cells were washed and incubated in DMEM for 30 min, and then infected with *S.* Typhimurium at MOI 100 for 30 min. Subsequently, media were replaced with 50 µg/ml gentamycin and 75 nM LysoTracker^TM^ DeepRed (L12492, Invitrogen) in DMEM. Cells were fixed with pre-warmed 4% PFA in PBS for 20 min at room temperature and mounted on glass slides with ProLong Diamond (P36971, Thermo Fisher Scientific).

### Microscope set-ups and image acquisition

Fluorescence images shown in Figs. 1, 2, 5 and 8 were acquired with a Zyla 4.2 sCMOS camera (Andor, Belfast, UK) and a spinning disk confocal module CSU-W1 (Yokogawa, Tokyo, Japan) mounted to a Nikon Ti2 eclipse microscope (Nikon, Düsseldorf, Germany). Images were acquired using a Plan Apo 20 x air/NA 0.75 objective, Plan Apo 60 x oil/NA 1.4 objective or Plan Fluor 100 x oil/NA 1.3 objective (all Nikon), 405/488/561/638 nm laser lines (Omicron, Rodgau-Dudenhofen, Germany) and appropriate filters controlled by NIS-Elements software (Nikon). Living cells were environmentally controlled with humidified 5% CO_2_ at 37 °C in an Okolab stage top incubation chamber (Okolab, Pozzuoli, Italy). Data shown in Fig. 5 were imaged by 3D structured illumination microscopy (SIM) using a Nikon Ti-Eclipse Nikon N-SIM E microscope and a CFI Apochromat TIRF 100 × Oil/NA 1.49 objective (Nikon). For image acquisition, NIS-Elements software controlled an Orca flash 4.0 LT sCMOS camera (Hamamatsu), a Piezo z drive (Mad city labs, Madison, WI, USA), a LU-N3-SIM 488/561/640 laser unit (Nikon) using the 488 nm laser or the 561 nm laser at 100% output, and a motorized N-SIM quad band filter with separate emission filters. Z-stacks were acquired using a step size of 200 nm. The reconstruction parameters IMC 2.11, HNS 0.37, OBS 0.07 were used for slice reconstructions (NIS-Elements, Nikon).

**Figure 1.**
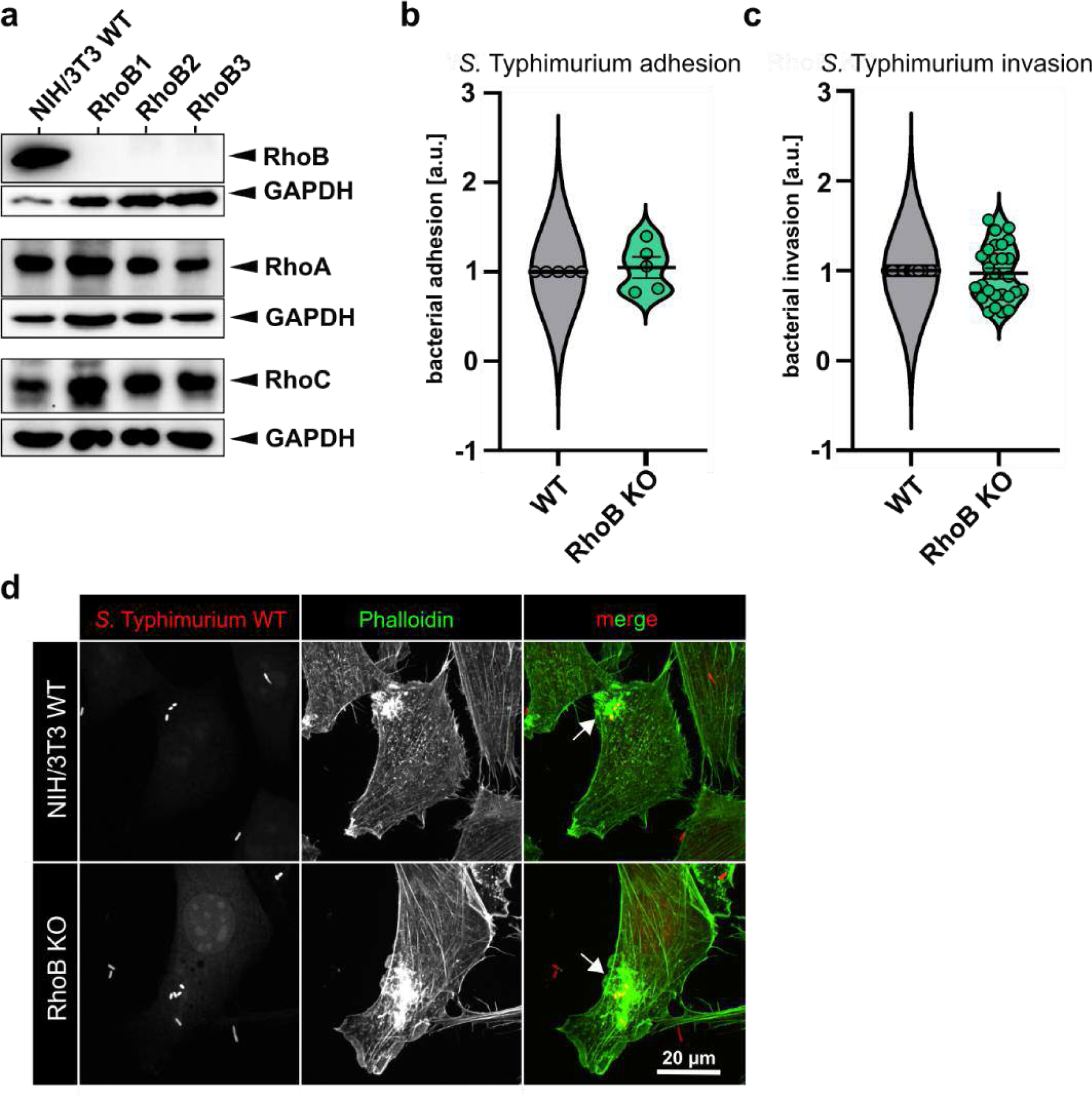
RhoB does not contribute to *S.* Typhimurium adhesion and invasion. **(a)** Western blots of individual RhoB clones in NIH/3T3 fibroblasts with RhoA, −B and −C specific antibodies, as indicated. GAPDH served as loading control. **(b, c)** NIH/3T3 WT and RhoB KO cells were assessed for *S.* Typhimurium WT adherence (15 min p.i.) **(b)** and invasion (60 min p.i.) **(c)**. Violin plots show quantifications of five independent adhesion and 28 independent invasion assays with at least three replicates each. Data are shown as means ± s.e.m. **(d)** NIH/3T3 WT and RhoB KO cells infected with wild-type *S.* Typhimurium for 15 min were stained with phalloidin. Images show maximum intensity projections of z-stacks from spinning disk microscopy. Arrows point to F-actin-rich ruffles induced by invasive *S.* Typhimurium.

**Figure 2.**
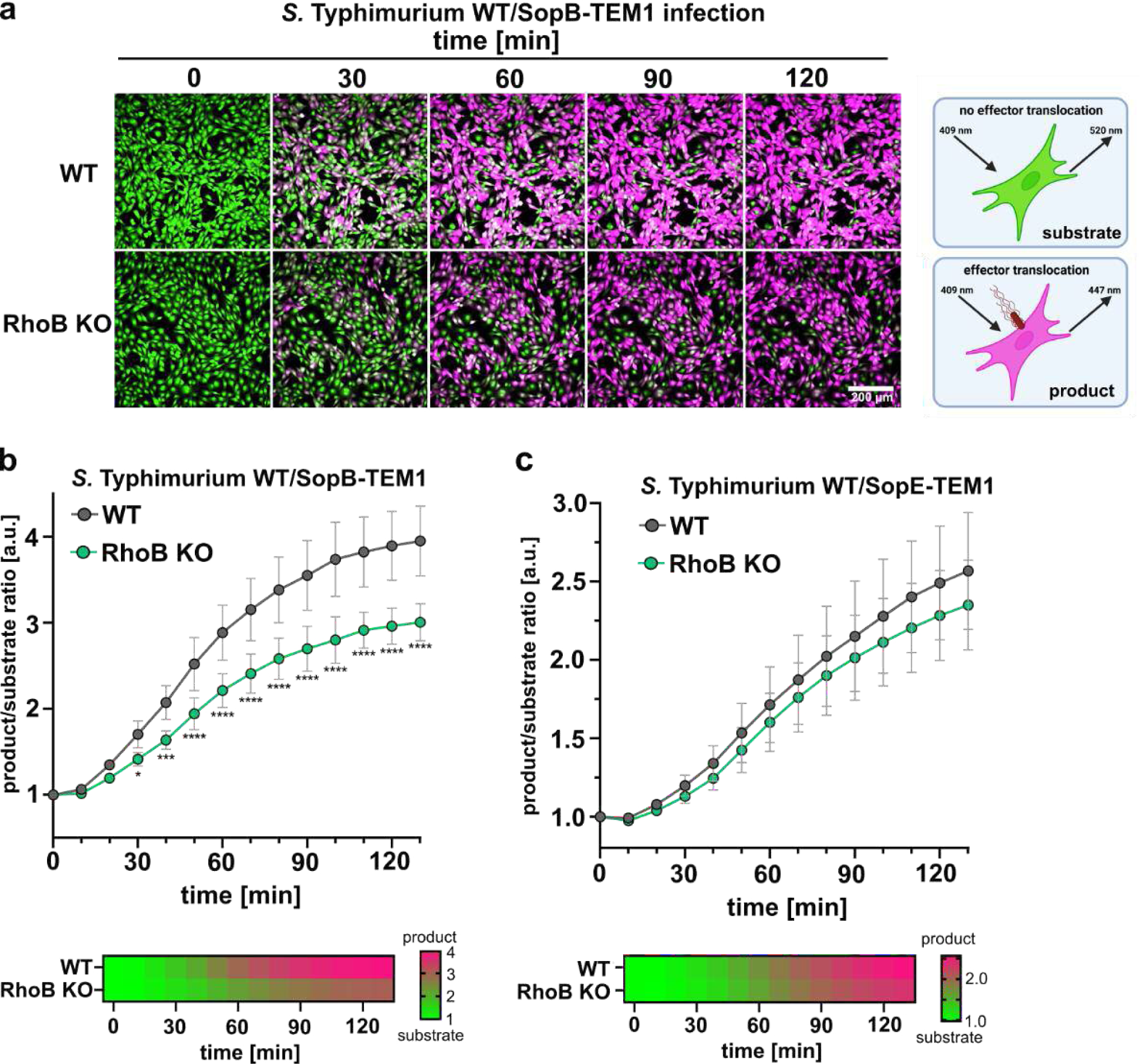
RhoB enhances effector secretion of SopB. **(a)** Cells loaded with the reporter dye CCF4-AM were infected with *S.* Typhimurium expressing SopB or SopE fused to β-lactamase (TEM1), as indicated. Upon effector translocation, the substrate CCF4-AM (pseudo-colored in green) is cleaved to the product CCF4 (pseudo-colored in pink). Translocation was monitored by spinning disk microscopy for 120 min. **(b, c)** Line graphs and heatmaps show fluorescence intensity quantification of the product over substrate ratio for SopB **(b)** and SopE **(c)** translocation in wild-type and RhoB KO cells, as indicated.

### Data Processing and Statistical Analyses

Images were analyzed using Fiji (https://imagej.net/software/fiji/), IncuCyte software (Sartorius, Göttingen, Germany) and NIS-Elements (Nikon). Further data processing and statistical analysis was carried out using NIS-Elements (Nikon), Fiji, Inkscape (Inkscape Project, https://inkscape.org), Prism 9 (Graphpad, San Diego, CA, USA) and Excel 2016 (Microsoft Corporation, Redmond, WA, USA). Details from statistical test, sample sizes and number of experiments are indicated in respective figure legends.

## Results

### RhoB does not contribute to *Salmonella* adhesion and invasion

RhoB has been reported to localize to *S.* Typhimurium invasion sites, however, RNAi-mediated knockdown of RhoB did not hamper invasion capacity of *S.* Typhimurium (Truong et al., 2018). Interestingly, RhoB localizes to endosomal vesicles (Fernandez-Borja et al., 2005; Mellor, Flynn, Nobes, Hall, & Parker, 1998), however, a potential function during later, intracellular stages of S. Typhimurium has remained unstudied. To learn about the role(s) of RhoB in these processes, we generated RhoB knockout cells by genome editing of the NIH/3T3 fibroblast cell line using CRISPR/Cas9. Several single cell-derived clones (RhoB1, RhoB2 and RhoB3) were isolated, genotyped by sequencing the targeted region of exon 1 and analyzed by Western Blotting for loss of the gene product (Fig. 1a, Supplementary Fig. S1a, b). In all of our RhoB knockout cell lines, no remaining protein was detectable, but we did not observe any reduction of expression of other Rho GTPase family members, i.e. RhoA and RhoC, as expected (Fig. 1a). For all types of experiments that are presented here, we have individually grown the three clones separately, and mixed them in equal numbers prior to each experiment, to exclude clonal differences. To first explore or exclude a potential contribution of RhoB in early steps of *S.* Typhimurium infection, we probed bacterial adhesion and invasion. We performed adhesion and gentamycin protection assays in NIH/3T3 WT and the RhoB KO population, and found that the lack of RhoB did neither affect *S.* Typhimurium adhesion nor its invasion capabilities (Fig. 1b, c), in line with a previous study that employed RNA interference to suppress RhoB expression (Truong et al., 2018). *S.* Typhimurium effectors activate a subset of host Rho-GTPases, including Cdc42 and Rac, leading to membrane ruffling (Hardt et al., 1998), which, however, is not an essential prerequisite for invasion (Hanisch et al., 2010). We next tested the appearance of invasion-triggered membrane ruffles through F-actin staining and fluorescence microscopy. As expected, cells lacking RhoB were still able to form membrane ruffles at *S.* Typhimurium invasion sites, virtually identical to their WT counterparts (Fig. 1d).

### RhoB enhances effector translocation of SopB

Since RhoB is recruited to the *Salmonella* invasion site at the plasma membrane, and since other Rho GTPases were described to contribute to T3SS-mediated effector translocation of *Yersinia pseudotuberculosis* (Mejia, Bliska, & Viboud, 2008; Schweer et al., 2013; Wolters et al., 2013), we next addressed the relevance of RhoB for *S.* Typhimurium effector secretion into the host cell. Translocation of the effector proteins SopB and SopE was monitored by time-lapse microscopy of *S.* Typhimurium expressing plasmid-encoded effector proteins fused to a reporter gene (TEM1) (Supplementary Fig. S2a). In fact, we found a significant decrease of SopB effector translocation efficiency in RhoB KO cells, detectable as early as 30 min pi (post-infection) and sustaining for more than 2 hpi (hours post infection) (Fig. 2a, b), while the translocation efficiency of SopE was only slightly reduced in RhoB KO cells (Fig. 2c, Supplementary Fig. S2b). The kinetic course of effector translocation thus revealed that in contrast to SopB, SopE is translocated largely independently of RhoB (Fig. 2c). Hence, our data reveal a role of RhoB in boosting SopB effector translocation through the T3SS.

### RhoB expression levels are upregulated in a SopB dependent manner

To better understand the role of RhoB during *S.* Typhimurium infections, we assessed RhoB expression levels at different time points of infection. Of note, RhoB expression levels are low under basal conditions and can be rapidly induced under cell stress conditions (Fritz, Kaina, & Aktories, 1995; J. Huelsenbeck et al., 2007; Prendergast, 2001; Zalcman et al., 1995; Zhang, Fu, Bu, Yao, & Wang, 2017). We first assessed the kinetics of RhoB expression levels during *S.* Typhimurium infection over a time period of 6 hpi. Indeed, RhoB protein levels were found to be significantly increased by approximately 2-to 3-fold at 2 hpi with wild-type *S.* Typhimurium. Strikingly, RhoB levels remained almost unaltered during infection with the mutant lacking SopB (ΔSopB) (Fig. 3a, b). When assessing the mRNA levels of RhoB during infection with wild-type *S.* Typhimurium, we detected a strong increase of expression in the first 1-2 hours, while the levels returned to the initial range at 4 hpi (Fig. 3c). From this we conclude that RhoB expression is induced upon infection by increased transcription, and that the resulting protein is then stabilized to sustain the protein level, as RhoB is known to have a remarkably short half-life of 30 min (Zalcman et al., 1995).

**Figure 3.**
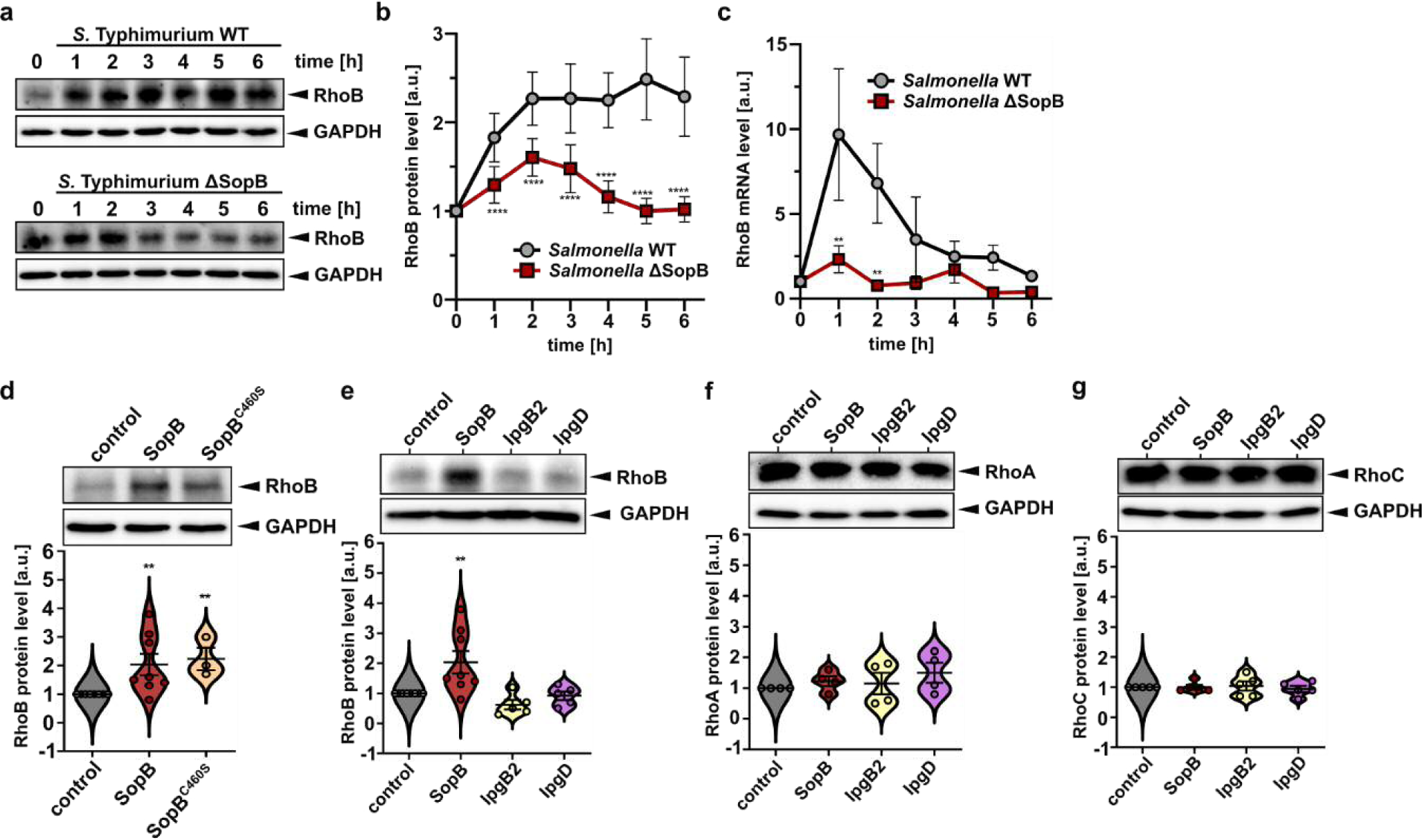
RhoB expression levels are upregulated in a SopB-dependent manner at later stages of *S.* Typhimurium infection. **(a,b)** RhoB expression levels were determined 0, 1, 2, 3, 4, 5, 6 hpi in *S.* Typhimurium wild-type (upper panel) or ΔSopB mutant (lower panel) infected NIH/3T3 wild-type cells by Western blotting. GAPDH served as loading control. **(b)** Quantifications of relative protein ratios of RhoB to GAPDH, as indicated. Data show means of protein levels derived from at least nine independent experiments ± s.e.m. ****p < 0.0001, two-way ANOVA. **(c)** Quantification of RhoB mRNA levels by RT-qPCR in NIH/3T3 wild-type cells infected with *S.* Typhimurium wild-type or ΔSopB mutant at different time points, as indicated. Expression values are relative to uninfected control. Data are means ± s.e.m. derived from three independent experiments. **p < 0.01, two-way ANOVA. **(d-g)** NIH/3T3 WT cells were transfected with SopB and the enzyme-dead mutant SopB^C460S^ or with plasmids expressing bacterial effector proteins SopB, IpgB2 or IpgD. RhoB **(d, e)**, RhoA **(f)** and RhoC **(g)** protein levels were assessed by Western blotting. GAPDH was used as loading control. Quantifications of at least three independent experiments are displayed as violin plots, with means ± s.e.m. **p < 0.01, two-way ANOVA.

We then asked if SopB alone, i.e. without translocation or in the absence of infection, is already sufficient to induce elevated RhoB expression and if so, if this requires the enzymatic PIPase activity of SopB. To explore this, we transfected NIH/3T3 WT cells with wild-type SopB and the enzyme dead mutant SopB^C460S^. Strikingly, the sole presence of SopB was sufficient to induce RhoB expression and both, SopB and the phosphatase inactive mutant SopB^C460S^ were capable of upregulating RhoB protein levels by approximately 2-fold (Fig. 3d). Finally, we compared this effect of SopB with that of the related effector proteins IpgD and IpgB2 from *Shigella flexneri*: IpgD is the orthologue of SopB, harboring C-terminal PIPase and N-terminal Cdc42-binding activities similar to SopB (Rodriguez-Escudero et al., 2011) and IpgB2, a GEF for RhoA (Klink et al., 2010) is potentially able to supplement for the RhoA activating effect of SopB (Hanisch et al., 2011). As a matter of fact, only SopB and SopB^C460S^ were able to elevate RhoB protein expression levels, but this was not observable for the *Shigella* effectors IpgD or IpgB2 (Fig. 3 d, e). To ensure that the effect is a GTPase-specific response, we probed whether these effectors, namely SopB, IpgB2 and IpgD affected expression levels of the closely related Rho GTPases, RhoA or RhoC. However, the expression levels of both were found to be unchanged for all effector proteins tested, confirming that *Salmonella* SopB specifically upregulates RhoB of the host (Fig. 3f, g). Collectively, RhoB expression is strongly upregulated upon infection with wild-type *S.* Typhimurium, which is dependent on the effector protein SopB, while RhoA and RhoC levels remain unaltered.

### RhoB localizes around intracellular bacteria and binds SopB

RhoB was described as a regulator of processes like actin reorganization, gene expression and vesicle trafficking (Vega & Ridley, 2018) and localizes to both the plasma membrane and endosomal vesicles (Adamson, Paterson, & Hall, 1992; Fernandez-Borja et al., 2005). Since RhoB was reported to localize to *S.* Typhimurium invasion sites (Truong et al., 2018), we wondered whether RhoB remains on the phagosomal membrane after invasion, as RhoB can also be found on other endosomal membranes. For this purpose, wild-type NIH/3T3 cells ectopically expressing EGFP-tagged RhoB were infected with *S.* Typhimurium. Cells were fixed at 1 hpi and RhoB was found to localize at the plasma membrane and also specifically accumulated around virtually all intracellular *bacteria* (Fig. 4a). However, at 6 hpi, RhoB localization to *S.* Typhimurium has disappeared (Fig. 4a lower panel). Together, our data uncover that RhoB is recruited to the plasma membrane by invading *S.* Typhimurium and remains associated with the early, immature SCV. To further assess the spatial association between these two proteins, we performed co-immunoprecipitation assays using NIH/3T3 cells co-transfected with EGFP-tagged RhoB and myc-tagged SopB. Strikingly, SopB specifically co-immunoprecipitated with RhoB while the unrelated control proteins, myc-tagged IRSp53 (Insulin receptor substrate 53) and EGFP did not (Fig. 4b). Hence, we here show for the first time an interaction of the *S.* Typhimurium effector SopB with RhoB. Moreover, RhoB contributes to secretion of the effector SopB, while SopB causes the significant upregulation of RhoB in infected cells, indicative of a positive feedback loop.

**Figure 4.**
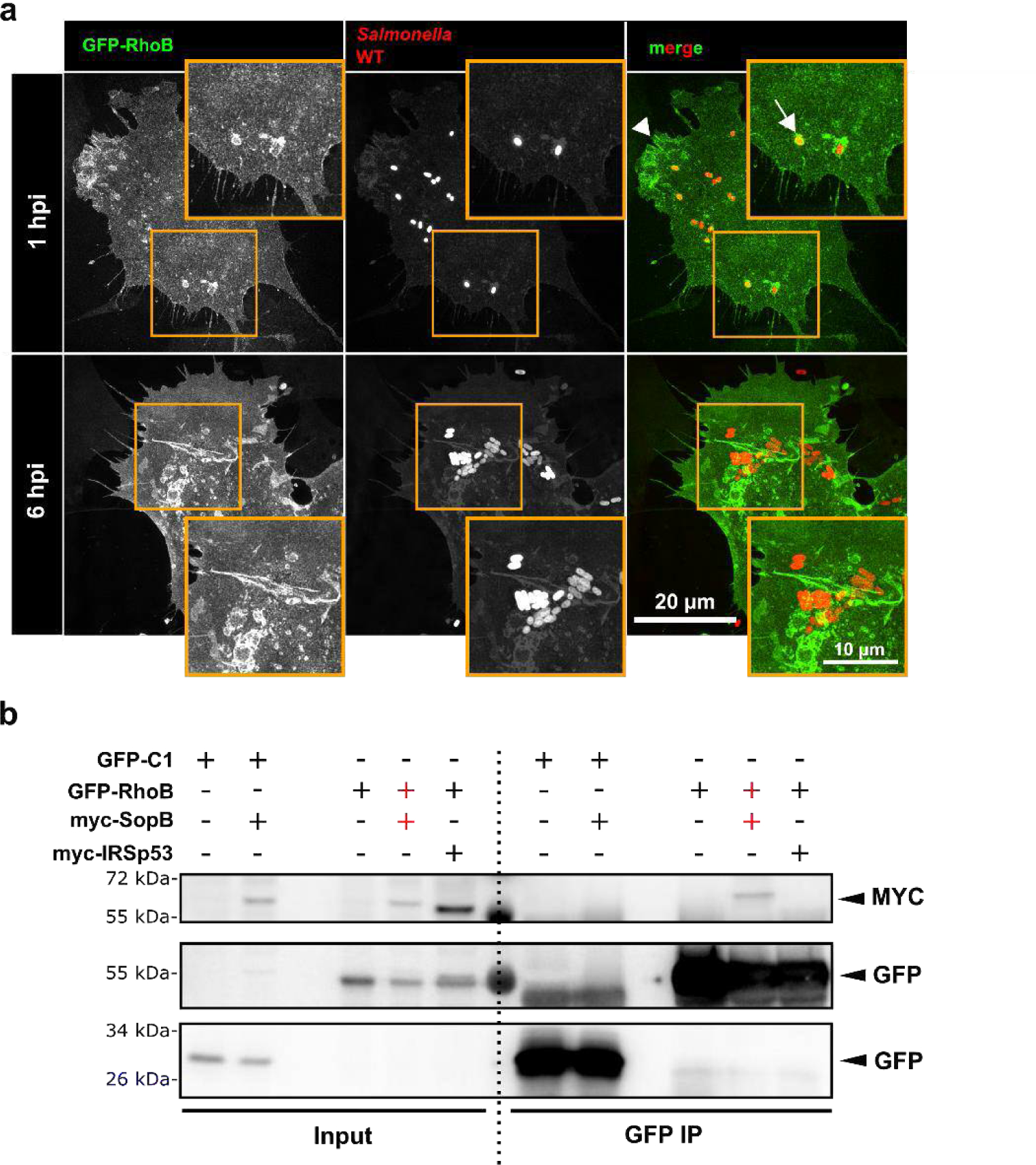
RhoB localizes around intracellular *Salmonella* and binds to SopB. **(a)** NIH/3T3 wild-type cells expressing EGFP-RhoB were infected with *S.* Typhimurium wild-type expressing mCherry. Cells were fixed 1 hpi and 6 hpi, stained with LysoTracker and imaged using superresolution microscopy (SIM). Images show maximum intensity projections. **(b)** NIH/3T3 wild-type cells were co-transfected with EGFP-C1, EGFP-RhoB, myc-SopB or myc-IRSp53. After 24 h, cell lysates were harvested and immunoprecipitation was performed with anti-GFP antibodies and analyzed by Western blotting. The anti-myc membrane was developed and then incubated with anti-GFP. n=3 independent experiments.

### RhoB contributes to *Salmonella* survival during later stages of infection

Since RhoB is not required for invasion itself, our findings prompted us to investigate the role of RhoB during later stages of infection. Importantly, gentamycin protection assays, used to measure the number of intracellular bacteria after different infection times, revealed that the survival of intracellular wild-type *S.* Typhimurium in cells lacking RhoB is significantly reduced, specifically by 45% and 60% after 6 and 12 hpi, respectively (Fig. 5a), suggesting that RhoB contributes to either replication or survival or both of intracellular *S.* Typhimurium.

**Figure 5.**
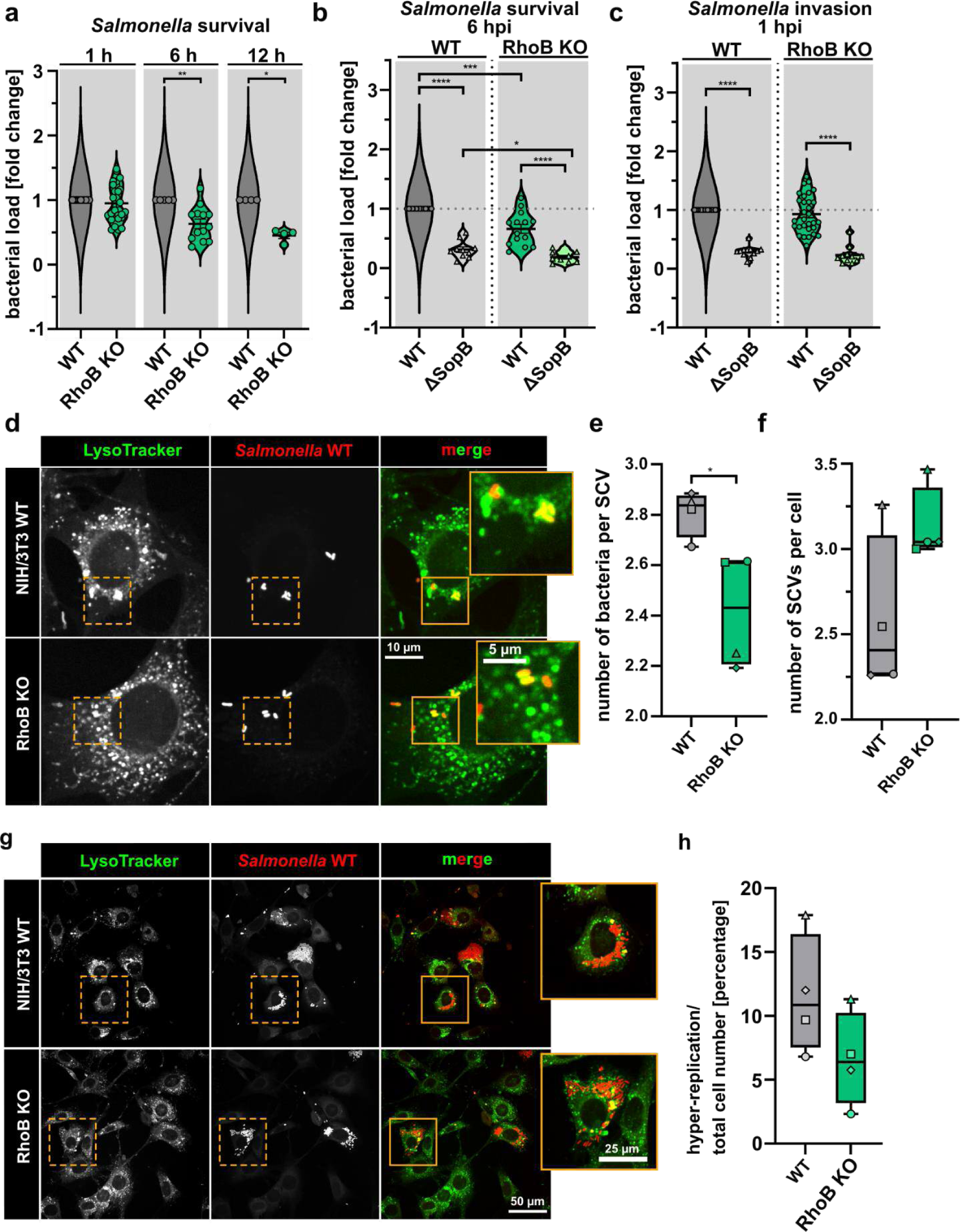
RhoB contributes to *Salmonella* survival during later stages of infection. **(a)** Relative numbers of wild-type *S.* Typhimurium in NIH/3T3 WT and RhoB KO cells at 1h, 6h and 12 hpi as assessed by gentamycin protection assays. Violin plots display means from at least four independent experiments with 5 replicates each ± s.e.m. * p < 0.05, ** p < 0.01, Mann-Whitney rank sum test. **(b, c)** Relative survival rates (6 hpi) **(b)** and invasion rates (1 hpi) **(c)** of *S.* Typhimurium wild-type and ΔSopB mutant in NIH/3T3 WT and RhoB KO cells, as indicated. Quantifications from at least twelve independent experiments with three replicates each are displayed as violin plots, with mean ± s.e.m. *p < 0.05, *** p < 0.001, **** p < 0.0001, Mann-Whitney rank sum test. **(d)** NIH/3T3 wild-type (upper panel) and RhoB KO (lower panel) cells were infected with mCherry*-*expressing wild-type *S.* Typhimurium for 6 h. Cells were stained with LysoTracker to visualize acidified vesicles such as lysosomes and imaged by spinning disk microscopy. Images show maximum intensity projections. **(e, f)** Quantification of number bacteria per SCV **(e)** and number of SCVs per infected cell **(f)**. The whiskers of the box and whiskers plots show the min to max of the data and the median. Data are from four independent experiments. * p < 0.05, Mann-Whitney rank sum test. **(e)** Hyper-replication events (≥30 bacteria per cell) of wild-type *S.* Typhimurium were assessed in NIH/3T3 wild-type and RhoB KO cells 6 hpi. **(h)** Quantification of cells with bacterial hyper-replication in NIH/3T3 wild-type and RhoB KO, as indicated. The whiskers of the box and whiskers plots show the min to max of four independent experiments and the median.

In order to learn if this phenomenon is connected to the interaction between SopB and RhoB, we performed a comparative analysis of the survival rates of WT and ΔSopB bacteria in WT *versus* RhoB KO cells. Intracellular survival rates of both, wild type and ΔSopB *S.* Typhimurium were statistically significantly decreased in RhoB KO cells (Fig. 5b, 6 hpi), but the reduction was most pronounced in the absence of both RhoB and SopB. Next, we asked to what extent the reduced number of ΔSopB bacteria may be due to reduced invasion, as it was shown before that ΔSopB bacteria invade much less effectively (Hanisch et al., 2011). Thus, we assessed the relevance of SopB for initial invasion into RhoB-deficient hosts: The degree of reduction caused by the lack of SopB was approximately 75% and virtually identical in wild-type and RhoB KO cells (Fig. 5c). Together, we conclude that RhoB is not involved in the initial invasion of *S.* Typhimurium, neither in the presence (Fig. 1b) nor in the absence of SopB (Fig. 5c), whereas it indeed plays a role in intracellular replication and/or survival of bacteria post invasion.

**Figure 6.**
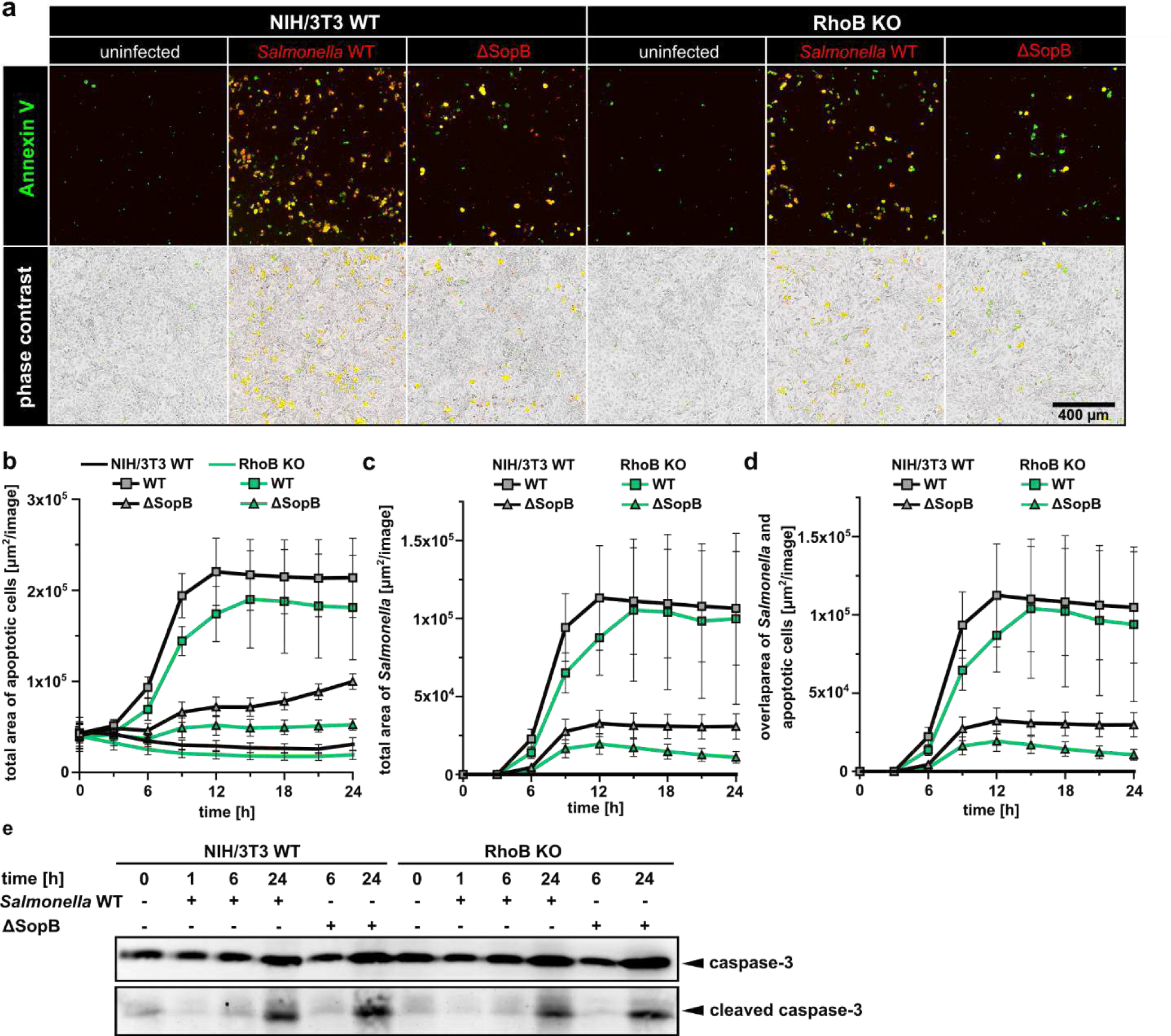
RhoB does not affect *Salmonella*-induced apoptosis. **(a)** NIH/3T3 and RhoB KO cells were infected with *S.* Typhimurium wild-type and ΔSopB mutant expressing mCherry, stained with annexin V Alexa Flour 488 and imaged by phase contrast and fluorescence microscopy over 24 hours. Images show respective overlays at 24 hpi. **(b-d)** Quantification of the total area of apoptotic cells per image **(b)**, total area of *S.* Typhimurium per image **(c)** and the overlapping area of apoptotic cells and *S.* Typhimurium **(d)**. Data show means from seven independent experiments with at least 4 replicates each ± s.e.m. **(e)** NIH/3T3 wild-type and RhoB KO cells infected with S. Typhimurium wild-type and ΔSopB were lysed in Laemmli buffer at indicated time points as indicated. Protein expression levels of caspase-3 and cleaved caspase-3 were assessed by Western blotting. Representative example of 3 independent experiments.

Since RhoB localizes to endomembranes regulating endocytic trafficking (Fernandez-Borja et al., 2005; Michaelson et al., 2001), we addressed whether the presence of RhoB might impact on the maturation and/or function of SCVs. Bacteria within SCVs were identified by co-localization of the bacteria with LysoTracker dye as described before (Marsman, Jordens, Kuijl, Janssen, & Neefjes, 2004). Of note, the SCV membrane closely envelopes the bacteria, leaving no visible space between the bacterial cell wall and the host-derived SCV membrane. Quantification of bacteria at 6 hpi within LysoTracker-positive compartments revealed that the number of bacteria per SCV is significantly decreased (Fig. 5d, e) while the number of SCVs per infected cell was slightly increased in RhoB KO cells (Fig. 5f). Hence, infected RhoB KO cells carry more and smaller SCVs with overall fewer bacteria. We furthermore observed that *S.* Typhimurium evade from the SCV and hyper-replicate in the host cytosol as described earlier (Knodler et al., 2014; Knodler et al., 2010; Malik-Kale et al., 2012). Nonetheless, *Salmonella* replication displays an obvious defect in RhoB KO cells. Thus, we assessed the differential contributions of SCV-encased *versus* hyper-replicated, cytosolic bacteria to the overall numbers of intracellular bacteria. Interestingly, *S.* Typhimurium displays reduced hyper-replication in cells lacking RhoB (Fig. 5g, h), explaining the majority of the difference in bacterial load between WT and RhoB KO cells. Although this experiment shows high variability in the number of hyper-replicative cells, the respective numbers were reduced for RhoB KO cells in every experiment (Fig. 5h).

### RhoB does not affect *Salmonella* induced-apoptosis

Inspired by the notion that reduced bacteria counts in RhoB KO cells are due to decreased cell numbers displaying hyper-replication of intracellular bacteria, we wondered if we lose bacteria due to apoptotic cell death specifically of infected cells in the absence of RhoB. In this case, bacteria would be lost as the extracellular medium in our setting contains non plasma membrane-permeable gentamycin to ensure that we only count intracellular bacteria. Apoptosis occurs as consequence of a pro-inflammatory response to *Salmonella* infection, eventually leading to programmed cell death of the host cell (Bertheloot, Latz, & Franklin, 2021). However, RhoB is also capable of contributing to AKT activation known to counteract apoptosis (A. Liu & Prendergast, 2000), which was presumed to also happen during *Salmonella* infection (Truong et al., 2018). In any case, *S.* Typhimurium infection is well established to induce apoptosis in host cells accompanied by activation of caspase-3 (Paesold, Guiney, Eckmann, & Kagnoff, 2002; Srikanth et al., 2010; Takaya et al., 2005). To clarify if the decreased bacterial load over the course of infection in RhoB KO cells is accompanied by increased numbers of apoptotic cells, we stained RhoB KO cells infected by RFP-tagged *Salmonella* with fluorescent annexin V, and measured the amount of apoptotic cells as area of fluorescent signal per image over a time period of 24 h. Apoptosis was indeed progressing during *S.* Typhimurium infection in both wild-type and RhoB KO cells starting at 6 hpi and reaching a plateau around 12-15 hpi (Fig. 6a, b). This was accompanied by a noticeable and successive increase in the number of RFP-tagged *S.* Typhimurium starting at 3 hpi (Fig. 6a, c). In contrast to our assumption, apoptosis in infected and non-infected RhoB KO is slightly lower than in wild-type cells (Fig. 6a, d), revealing that increased apoptosis cannot explain reduced bacterial load in RhoB KO cells. We further solidified this conclusion by assessing protein levels of cleaved and non-cleaved caspase-3 by western blotting in *S.* Typhimurium infected WT *versus* RhoB KO cells. In line with our previous results, RhoB KO and wild-type cells displayed comparable levels of caspase-3 cleavage (Fig. 6e). This notion is further supported by the finding that AKT activation upon infection with *S.* Typhimurium is virtually identical in WT and RhoB KO cells (Supplementary Fig. S3). Therefore, neither induction nor suppression of apoptosis by *S.* Typhimurium depends on RhoB. Together with our previous findings (Fig. 4, Fig. 5), this suggests that the presence of RhoB on the early SCV membrane in the first 1-2 hours post invasion directly relates to bacterial survival (not host survival) and is either crucial to induce efficient replication and/or for later escape of the bacteria from the SCV.

### *Salmonella* induces autophagy in a RhoB-dependent mechanism

In *S.* Typhimurium infection, xenophagy has been shown to play a role in the host’s immune response to the bacteria (Wang, Yan, Niu, Huang, & Wu, 2018). Whereas it is in the interest of invaded bacteria to suppress this host response to aid persistence (Huang & Brumell, 2014), they might at the same time benefit from the mobilization of nutrients through housekeeping autophagy. In order to clarify if autophagy is involved in RhoB-mediated *S.* Typhimurium survival, we performed a series of analyses: First, we compared the autophagic flux in wild-type and RhoB KO cells. LC3-II and p62 levels were assessed in cells with both genotypes over a time course of 4 h in amino acid-deprived medium (HBSS) in the presence or absence of bafilomycin A1 (BafA), an inhibitor of the lysosomal vacuolar H^+^-ATPase (v-ATPase), blocking acidification of the lysosome and thus autophagosome-lysosome fusion (Klionsky, Elazar, Seglen, & Rubinsztein, 2008) (Fig. 7a, b). The lipidation of cytosolic LC3 (LC3-I) leads to the membrane-bound form (LC3-II) and characterizes autophagy induction (Klionsky et al., 2016). The autophagy marker p62/SQSTM1 links ubiquitinated cargo to the autophagic machinery and is degraded together with the cargo upon fusion with lysosomes (Bjorkoy et al., 2009). Autophagic flux upon nutrient deprivation was detectable in both, wild-type and RhoB KO cells, as revealed by LC3 conversion and p62 levels (Fig. 7a). Moreover, LC3-II accumulation was equally observed in wild-type and RhoB KO cells upon BafA1 treatment (Fig. 7b). However, whereas the ubiquitin adapter p62 accumulated in wild-type cells as expected, its levels remained low in RhoB KO cells over 4 hours following BafA1 treatment (Fig. 7b). From this, we conclude that nutrient deprivation can, in principle, initiate autophagy in RhoB KO cells, as judged by virtually identical levels of LC3 lipidation, while the degradation of ubiquitinated cargo appears altered.

**Figure 7.**
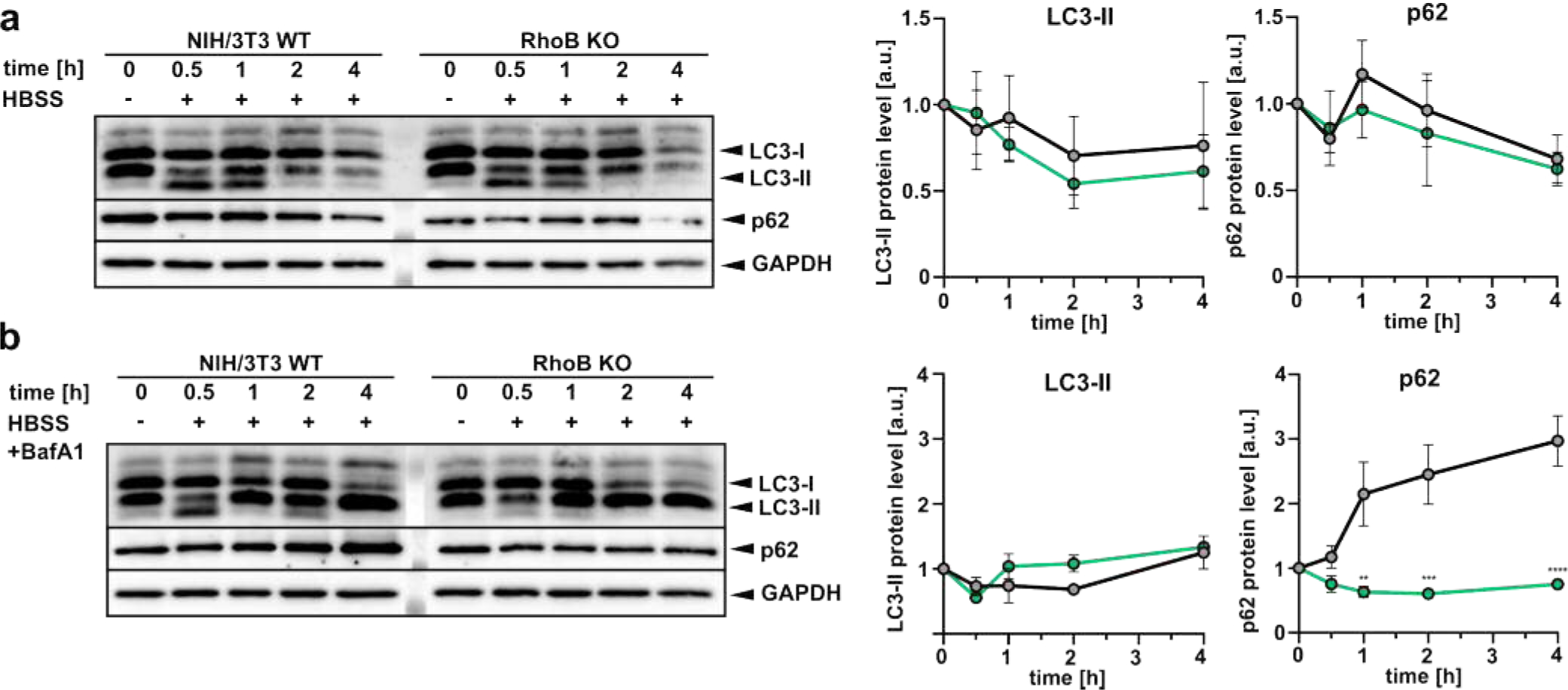
RhoB is dispensable for induction of autophagy by nutrient deprivation. **(a, b)** NIH/3T3 wild-type and RhoB KO cells were treated with HBSS in the presence **(b)** or absence **(a)** of 250 nM bafilomycin A1, and lysed in Laemmli buffer at the indicated time points. Protein levels of LC3 and p62 were assessed by western blotting. GAPDH served as loading control. Representative images (left), and line graphs on the right show quantification of LC3-II and p62 protein levels derived from three independent experiments, means ± s.e.m.

We next examined autophagy in the context of long-term *S.* Typhimurium infections: LC3-II was strongly increased in wild-type cells at 24 hpi (approx. 3-to 4-fold), which was similar for infections with both WT and ΔSopB bacteria (Fig. 8a, b). Strikingly however, induction of LC3 lipidation upon *S.* Typhimurium infection was completely abrogated in RhoB KO cells (Fig. 8a, b). To monitor autophagy induction during early stages of *S.* Typhimurium infection, samples of infected, serum-starved wild-type and RhoB KO cells were collected between 0-4 hpi, immunoblotted with LC3 and p62 antibodies and quantified (Fig. 8c). Whereas LC3-II and p62 levels strongly increased over time in wild-type cells, both LC3-II and p62 remained unaltered upon infection with *S.* Typhimurium in RhoB KO cells (Fig. 8c). This was also observed in ΔSopB infected cells (Supplementary Fig. S4).

**Figure 8.**
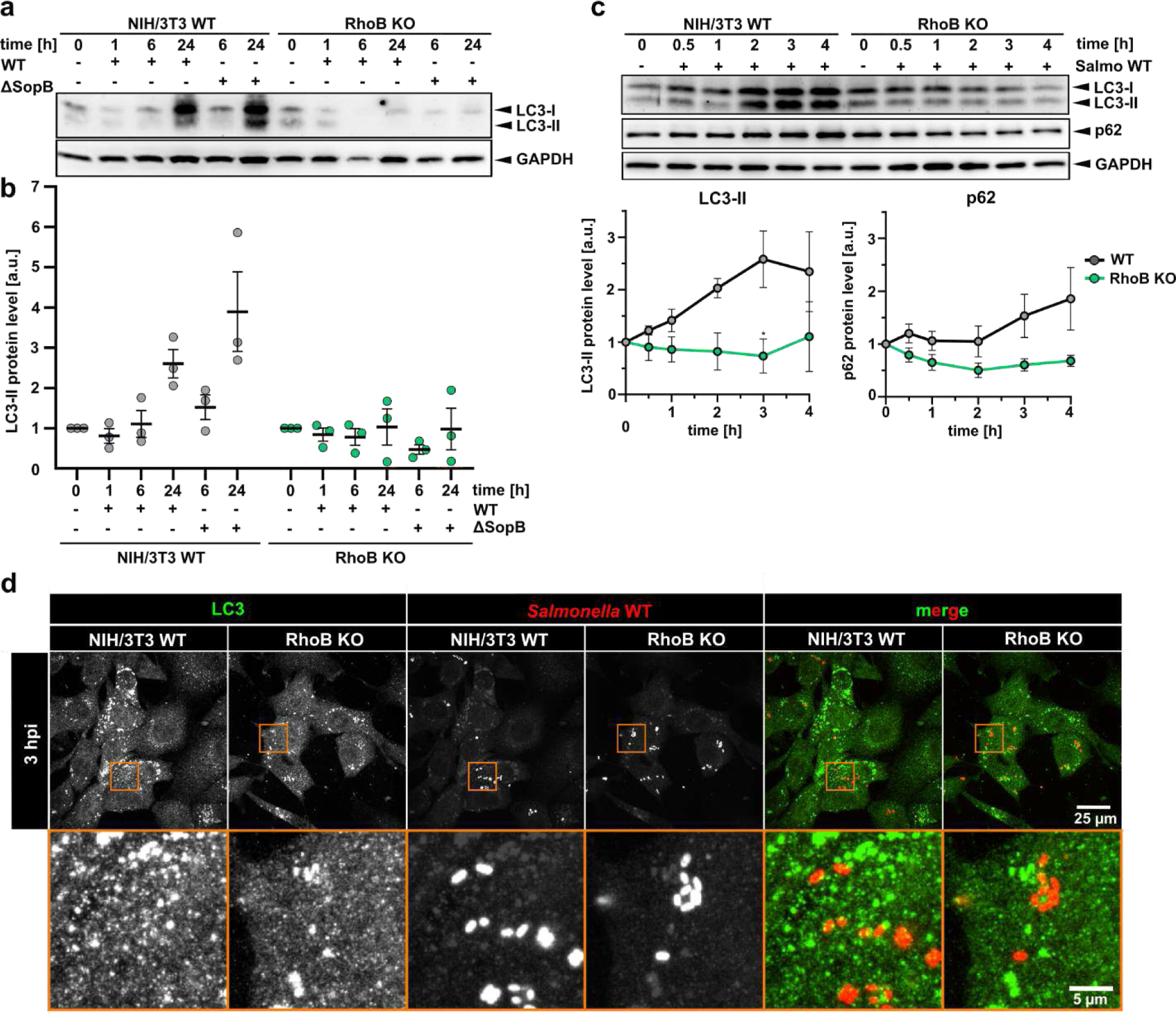
*Salmonella* induces autophagy in a RhoB-dependent mechanism. **(a, b)** NIH/3T3 wild-type and RhoB KO cells infected with *S.* Typhimurium wild-type or ΔSopB mutant were lysed in Laemmli buffer at indicated time points. Protein expression levels of LC3, p62 and GAPDH were assessed by Western blotting (representative blot). **(b)** Data show means of LC3-II protein levels derived from three independent experiments ± s.e.m. **(c)** Serum-starved wild-type and RhoB KO cells were infected with wild-type *S.* Typhimurium and lysed with Laemmli buffer at 0.5, 1, 2, 3, 4 hpi. LC3, p62 and GAPDH were assessed by Western blotting. Bottom panel: Quantifications of relative ratios of LC3 and p62 to GAPDH as indicated. Data are means of protein levels from three independent experiments ± s.e.m. * p < 0.05, two-way ANOVA. **(d)** NIH/3T3 wild-type and RhoB KO cells were infected with *S.* Typhimurium wild-type expressing mCherry. Cells were fixed at 1 and 3 hpi and immuno-stained for LC3 and bacteria. Images were acquired by spinning disk microscopy and show maximum intensity projections.

We next probed whether Salmonella-induced autophagy also leads to a co-localization of the autophagic marker LC3 with invading bacteria. Moreover, we interrogated if there is a difference in RhoB KO cells concerning LC3 localization. Indirect immunofluorescence with LC3-specific antibodies revealed that LC3 accumulations most likely representing autophagosomes were increased as compared to non-infected cells (Supplementary Fig. S5) in WT NIH/3T3, but less so in RhoB KO cells. More importantly, neither in WT-nor in RhoB KO cells we detected any obvious encasing of *Salmonella* by LC3, although LC3-positive vesicles were occasionally found in close vicinity to the SCV in both cell types (Fig. 8d).

Therefore, autophagy induction upon either nutrient deprivation or *S.* Typhimurium infection has fundamentally different effects in the absence of RhoB. The former (nutrient deprivation-induced) is initiated normally, whereas the latter (*Salmonella*-induced) is abrogated at the early stage of LC3 lipidation. Consequently, the initiation of *Salmonella*-induced LC3 conversion coincides with the localization of RhoB on the *Salmonella*-containing phagosome, where it likely co-localizes and interacts with SopB. Together, RhoB thus operates in early *Salmonella*-induced autophagy, aiding subsequent bacterial survival and hyper-replication.

## Discussion

Rho GTPases are molecular switches that quickly integrate signals and thereby regulate cellular processes such as actin-mediated cell motility, cell cycle progression or vesicle transport (Heasman & Ridley, 2008). During evolution, bacteria have been developing specific pathogenicity factors such as T3SS and effector proteins known to hijack host cell Rho GTPases in order to promote bacterial adhesion, invasion and persistence (Lemichez, 2017; Popoff, 2014). Whereas the roles of Rac1, Cdc42 and RhoA for *Salmonella* invasion have been explored, the role of other Rho GTPases such as RhoB during *S.* Typhimurium infection remained poorly understood. Strikingly, we here uncover for the first time that host cell RhoB contributes to *S.* Typhimurium survival in infected cells and reveal a so-far unknown functional interaction between the host factor RhoB and the *Salmonella* effector SopB.

Invasion of *S.* Typhimurium into non-phagocytic cells is mediated by the effector proteins SopB and SopE/E2 targeting host cell Rho GTPases RhoA and Rac1 or Cdc42, respectively. Translocation of these effectors is controlled by a T3SS and requires contact with the host cell (Lou, Zhang, Piao, & Wang, 2019; Lunelli, Hurwitz, Lambers, & Kolbe, 2011). Our work uncovers that T3 secretion leads to local recruitment of RhoB, which is then utilized by *S.* Typhimurium to enhance effector translocation of SopB into the host cell cytoplasm. Moreover, the presence of SopB in the host’s cytosol is sufficient to induce profound upregulation of RhoB expression. Thus SopB is (i) secreted in the first 2 hpi in a cumulative fashion, in line with a previous report (Patel, Hueffer, Lam, & Galan, 2009), (ii) its secretion is considerably promoted through host cell RhoB and (iii) RhoB is significantly upregulated by the sole presence of SopB in the host. This represents a canonical positive feedback between bacterial SopB and host cell RhoB, which is started by initial Type 3 secretion of SopB and continues for the first 1-2 hours following bacteria-host contact, again coincident with initial localization of RhoB at the invasion site (Truong et al., 2018) and later in infection on *Salmonella*-containing phagosomes. This is additionally accompanied by interaction of the two proteins. This effect of RhoB is specific to SopB translocation, and does not apply to SopE secretion, an effector that is secreted during very early stages of invasion (approx. 15 min pi) (Kubori & Galan, 2003).

Although our data suggest that local recruitment of RhoB to the site of Type 3 secretion regulates effector secretion of SopB, the molecular mechanism of how this is achieved remains elusive. Notably, however, CNF-Y has likewise been shown to enhance Yop translocation of *Yersinia pseudotuberculosis* by activating Rho GTPases, indicating that feedback between Rho GTPases and effective functioning of T3SSs might represent a common theme (Mejia et al., 2008; Schweer et al., 2013; Wolters et al., 2013).

Previous studies have shown that cells under basal conditions express low levels of RhoB and, in addition, that the half-life of the protein is relatively short (Fritz et al., 1995; Vega & Ridley, 2018). Stimuli like growth factors, cell cycle progression and cytokines, but also stressors such as UV irradiation or bacterial toxins like cytotoxic necrotizing factor 1 (CNF1) from *Escherichia coli* can rapidly lead to elevated RhoB expression (Gerhard et al., 2005; Gutierrez et al., 2019; J. Huelsenbeck et al., 2007; S. C. Huelsenbeck et al., 2013; Vega & Ridley, 2018). We unveil that SopB also shares this effect on RhoB expression and, noteworthy, it does so independently of its phosphatase activity. Moreover, this effect cannot be induced by expression of the SopB orthologue IpgD from *Shigella flexneri,* underscoring that this is a *Salmonella*-specific trait and likely relating to the intracellular life-style of *Salmonella* with the necessity to manipulate the phagosomal membrane. Although we do not understand the precise molecular mechanism yet, we delineate that RhoB expression is induced in at least two ways: (i) at the transcriptional level by short term upregulation of mRNA expression and (ii) by stabilizing the protein and protecting it from otherwise rapid degradation.

To clarify why *S.* Typhimurium benefit from RhoB upregulation, we found that *Salmonella* strongly induces a subtype of autophagy upon infection, a phenomenon that is strongly abrogated in the absence of RhoB. *S.* Typhimurium has developed strategies to replicate and persist intracellularly by formation of the SCV followed by phagosomal escape into the cytosol and hyper-replication (Knodler et al., 2010; Malik-Kale et al., 2012). Establishment of the SCV requires the effector-mediated recruitment of endosomal/lysosomal membranes to the phagosome (Bakowski, Braun, & Brumell, 2008; Drecktrah et al., 2007). One important pathogenic factor in SCV formation is SopB. It was shown to mediate host cell signaling, activation of Rho GTPases, activation of protein kinase B (AKT), fission of the SCV from the plasma membrane, recruitment of Rab5, Vps34, sorting nexins-1 and −3 (Braun et al., 2010; Bujny et al., 2008; Mallo et al., 2008) and perinuclear SCV positioning by myosin II activation (Wasylnka et al., 2008). Furthermore, SopB was reported to modulate surface charge of the SCV to avoid fusion with the lysosome (Bakowski et al., 2010). Our work uncovered RhoB as a novel host cell factor promoting intracellular *Salmonella* survival at later stages of infection. We also observed that RhoB is enriched around internalized *S.* Typhimurium at 1 hpi, but is lost from these sites at later stages such as at 6 hpi. The intracellular lifestyle of *S.* Typhimurium has recently been shown to be bimodal (Malik-Kale et al., 2012), with two populations of intracellular *Salmonella,* one that replicates within the SCV, which is SPI-2 dependent, and a SPI2-independent group replicating in the cytosol.

Both cytosolic and SCV-encased *Salmonella* were described to be targeted by the autophagic machinery (Wu, Shen, Zhang, Xiao, & Shi, 2020). *S.* Typhimurium infection in wild-type cells increases levels of LC3 lipidation, which is in line with earlier reports (W. Liu et al., 2018; W. Liu et al., 2019). Surprisingly, we found that *S.* Typhimurium-induced LC3 lipidation is completely abrogated in cells lacking RhoB. Moreover, this phenomenon is specific for infection, as serum-starvation revealed equal levels of autophagy induction in the presence and absence of RhoB. Our results thus uncover that *S.* Typhimurium usurps RhoB for the induction of a specific subtype of autophagy, which is in essence connected to intracellular survival of the bacteria. Finally, *Salmonella*-induced LC3-II does not localize to the SCV surrounding the bacteria, speaking against it being an antibacterial response of the host. The bulk of LC3-II-positive autophagosomes is not associated with the intracellular bacteria and only some autophagosomes are observed in the vicinity of SCVs. It is tempting to speculate therefore that *Salmonella* actively induces this type of autophagy perhaps in order to mobilize nutrients, and that a lack of *Salmonella*-induced autophagy in RhoB KO cells explains for the replicative disadvantage of bacteria in the absence of RhoB. In the absence of bacterial SopB, *Salmonella* do not only invade at reduced rates (Hanisch et al., 2011), but they also lack the positive feedback loop to induce RhoB, also resulting in a mild survival defect.

From this we conclude a model (Fig. 9), in which *Salmonella* actively induces autophagy to mobilize nutrients thus promoting long term survival and extra-vacuolar hyper-replication. This process absolutely requires the host cell factor RhoB, which is induced by the bacterial factor SopB in the first hpi via a positive feedback loop that is accompanied by co-localization and interaction of bacterial SopB and host RhoB.

**Figure 9.**
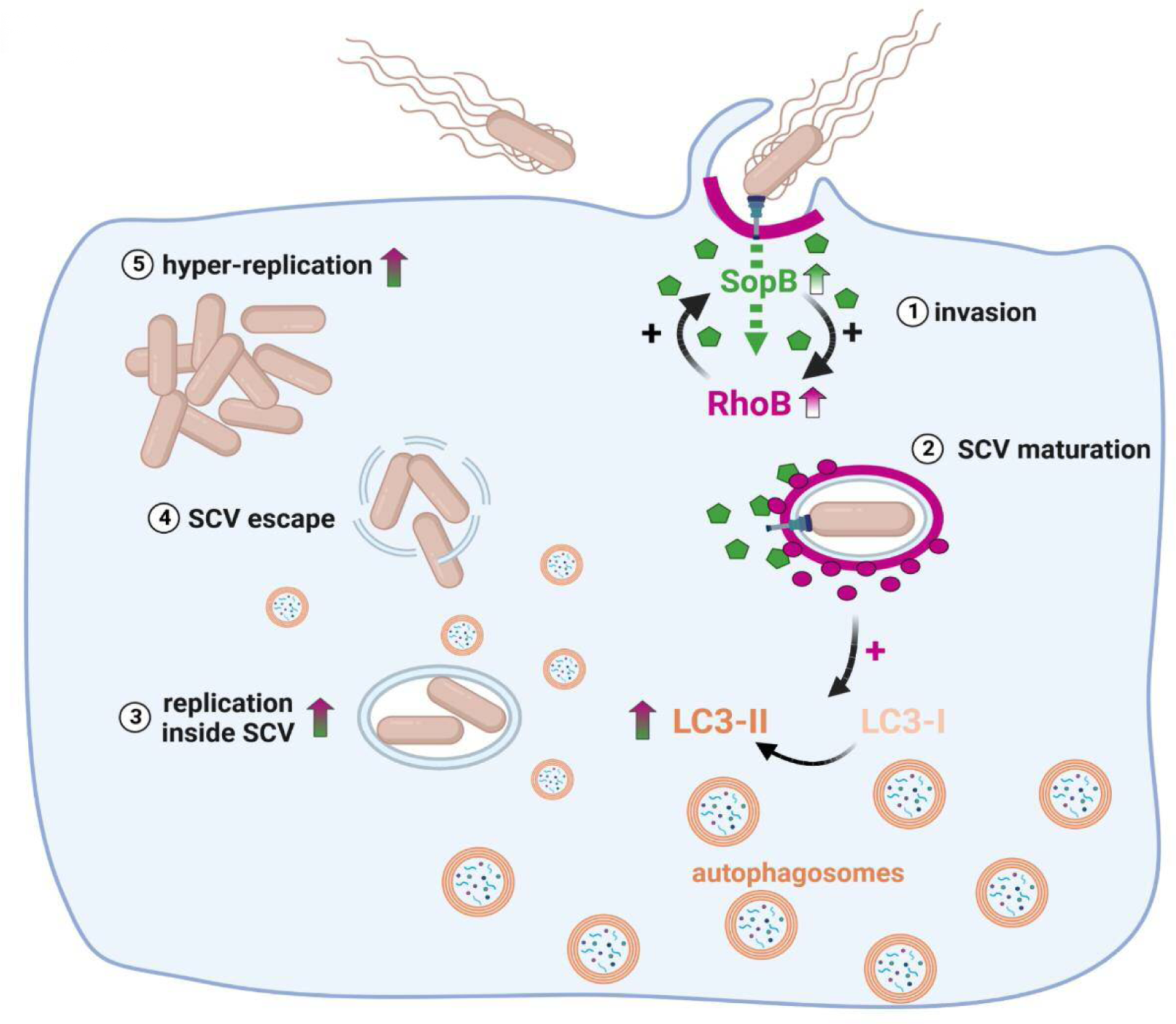
Roles of SopB and RhoB during *S.* Typhimurium invasion and replication. (1) *S.* Typhimurium translocates the effector protein SopB, which upregulates RhoB protein levels. Note that the presence of RhoB stimulates further SopB translocation, suggesting a positive feedback loop. (2) RhoB localizes to the maturing SCV and (3) eventually disappears. Moreover, *Salmonella* invasion specifically induces autophagy, a process that is completely abrogated in the absence of RhoB. In addition, RhoB supports both, replication of *Salmonella* inside the SCV (3) as well as hyper-replication in the host cell cytosol (5) after escape of *Salmonella* from the SCV (4).

The orthologous effector from *Shigella flexneri*, IpgD, which shares the domain composition with SopB and overall displays high levels of homology (Tran Van Nhieu, Latour-Lambert, & Enninga, 2022), cannot induce RhoB expression. This can be explained by the different survival strategies of *Salmonella versus Shigella*, with the latter escaping the phagosome right after invasion and then evading the antibacterial response by assembling host cell F-actin (Mostowy et al., 2010). Future research is needed to understand the molecular mechanisms of RhoB induction by SopB and the differences between *Salmonella*-induced as compared to starvation-induced autophagy. Notwithstanding this, our results establish yet another intriguing example of host-pathogen interaction, which highly specifically accommodates the needs of the pathogen, in this case intracellular survival and replication of *Salmonella* Typhimurium.

## Acknowledgements

We are grateful to Lothar Gröbe for flow cytometry and FACS sorting. This work was supported by the Deutsche Forschungsgemeinschaft (to KR, individual grant RO 2414/8-1) and by the HGF impulse fund (to TEBS, grant W2/W3-066).

## Author contributions

**Marco Kirchenwitz**: Conceptualization, Methodology, Validation, Formal analysis, Investigation, Writing - Original Draft preparation, Writing - Review & Editing, Visualization; **Jessica Halfen**: Investigation, Formal analysis; **Kristin von Peinen**: Investigation; **Silvia Prettin**: Investigation; **Jana Kollasser**: Investigation; **Cord Brakebusch**: Resources; **Klemens Rottner**: Writing - Review & Editing, Funding acquisition; **Anika Steffen**: Conceptualization, Validation, Writing - Review & Editing, Visualization, Supervision; **Theresia E.B. Stradal**: Conceptualization, Validation, Resources, Writing - Review & Editing, Visualization, Supervision, Funding acquisition

---

**Supplementary Figure S1.**
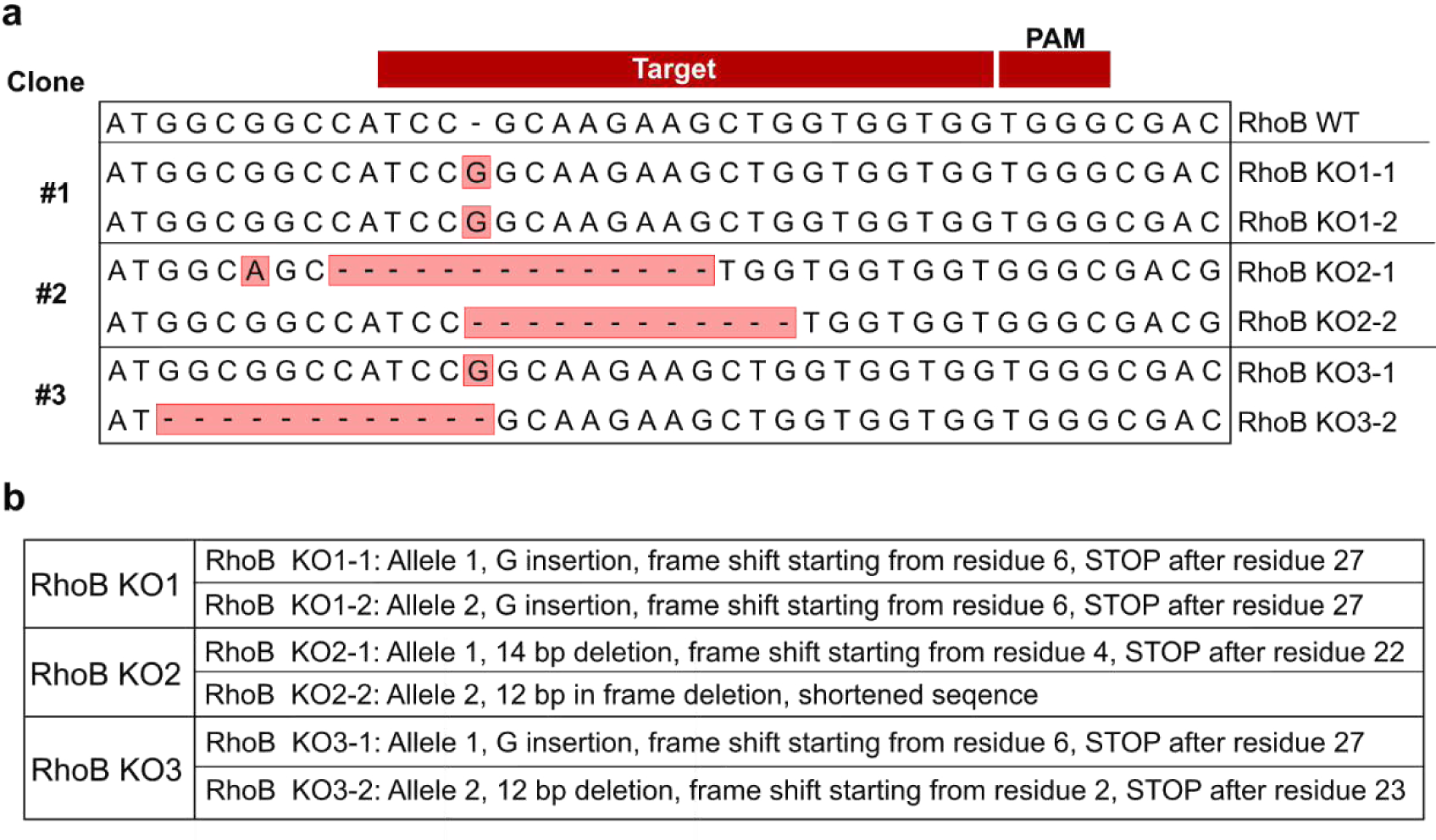
Sequencing results used to determine the RhoB knockout genotypes used in this study. (a) Sequencing results of mutated alleles in clones KO1, KO2 and KO3 are given. (b) Type of nucleic acid mutation and corresponding translation consequences are listed.

**Supplementary Figure S2.**
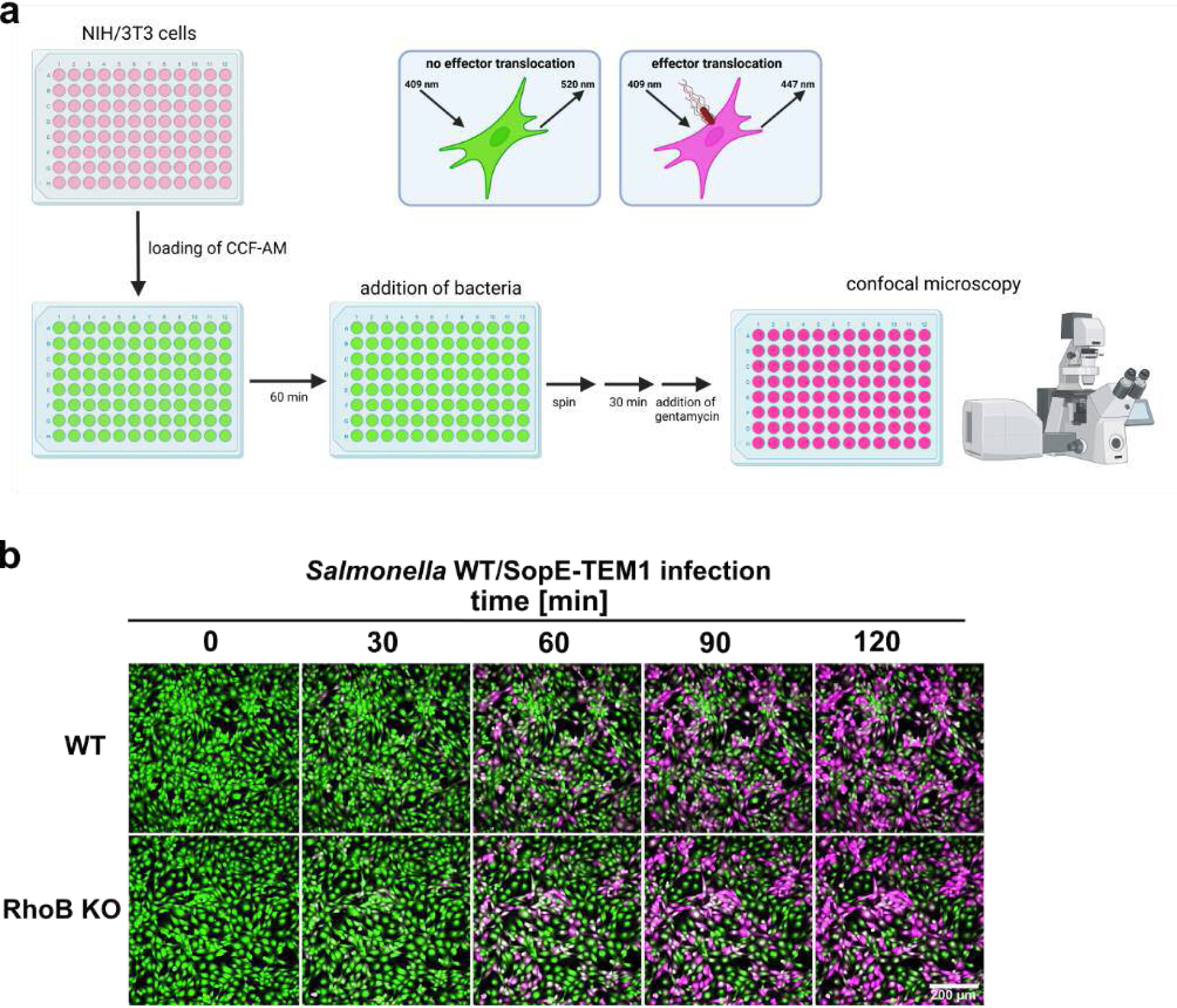
*Salmonella* effector translocation of SopE. **(a)** Experimental workflow of *S.* Typhimurium effector translocation assay in NIH/3T3 wild-type and RhoB KO cells. Images were created with BioRender.com. **(b)** Cells loaded with the reporter dye CCF4-AM were infected with *S.* Typhimurium expressing SopE fused to β-lactamase (TEM1). Upon effector translocation, the substrate CCF4-AM (green fluorescence emission at 520 nm) is cleaved to the product CCF4 (pseudo-colored in pink; fluorescence emission at 460 nm). Translocation was monitored by spinning disk microscopy up to 120 min, as indicated. No major differences could be discerned for SopE translocation in WT *versus* RhoB KO cells.

**Supplementary Figure S3.**
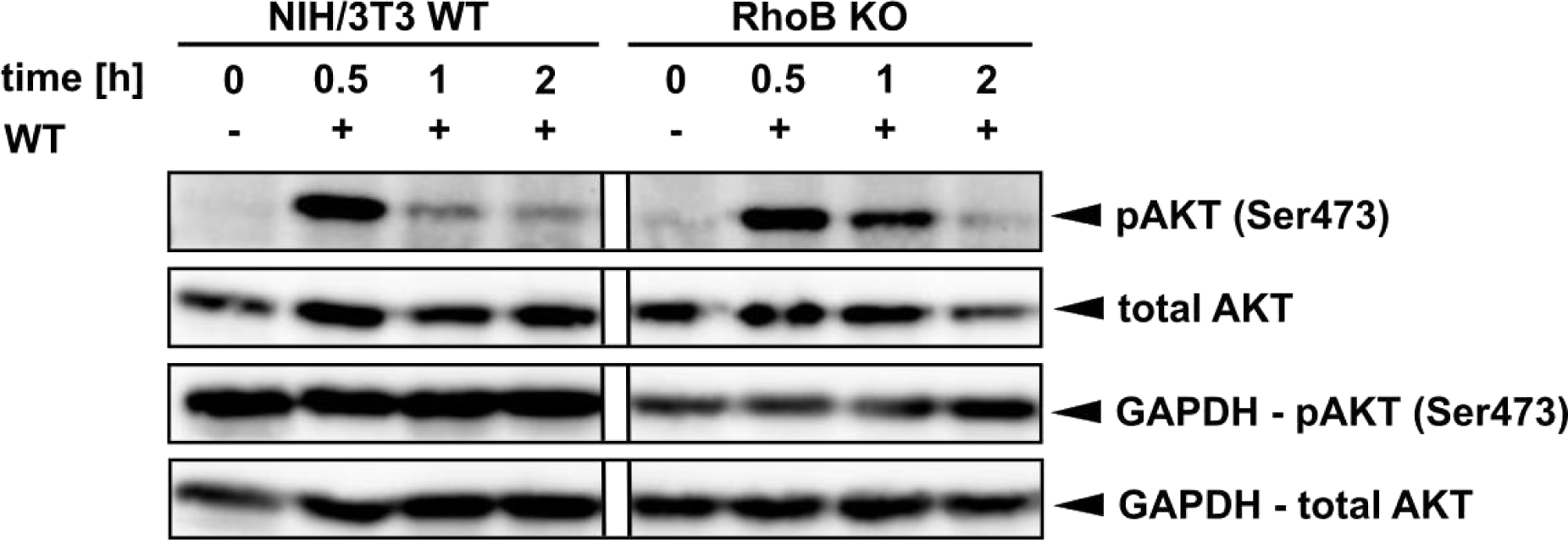
*Salmonella*-induced AKT activation is independent of host cell RhoB. NIH/3T3 wild-type and RhoB KO cells infected with *S.* Typhimurium wild-type were lysed in Laemmli buffer at time points as indicated. Protein expression levels of total AKT, phospho-AKT (Ser473) and GAPDH were assessed by Western blotting.

**Supplementary Figure S4.**
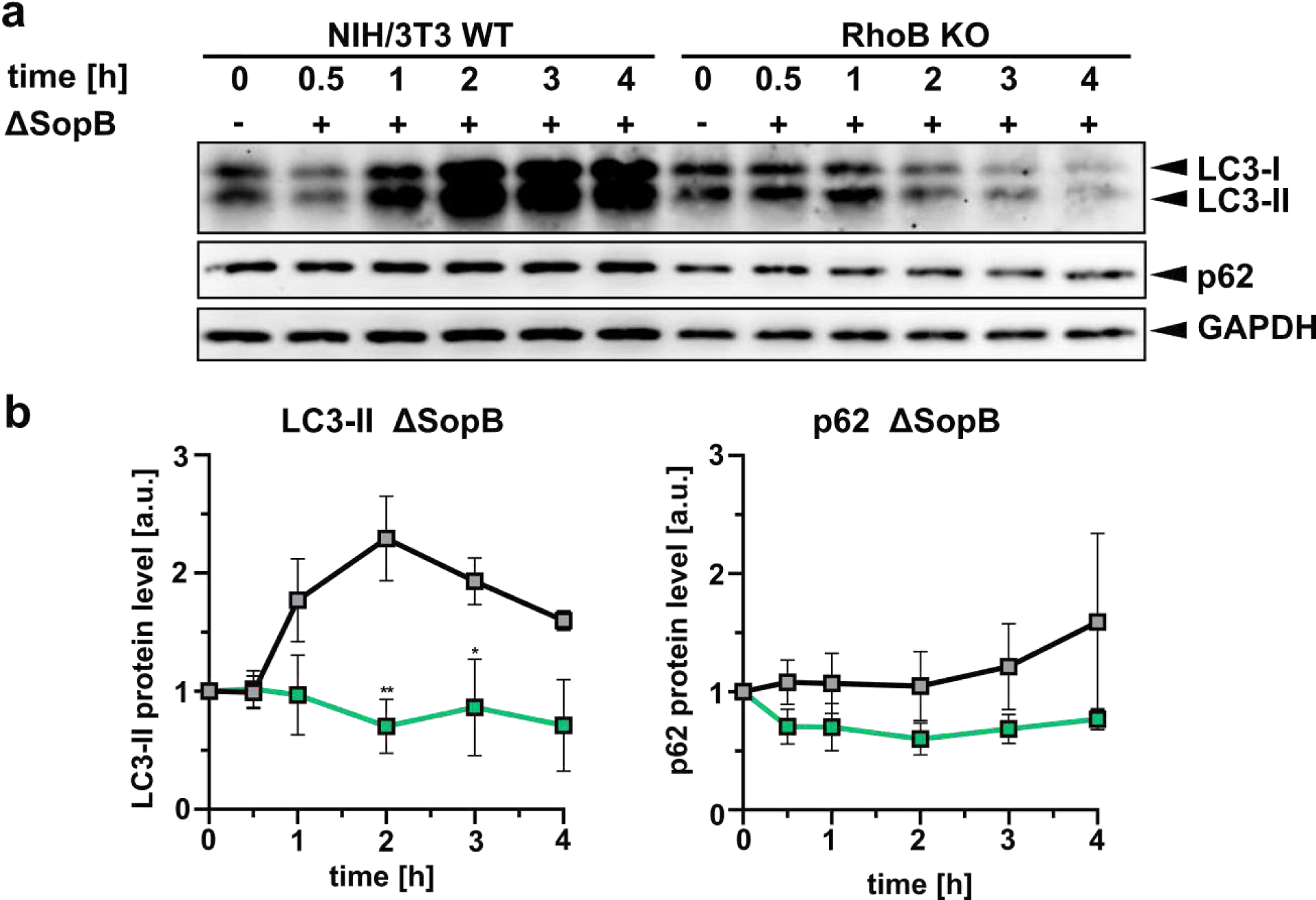
*Salmonella*-induced autophagy is independent of *Salmonella* SopB, but depends on host cell RhoB. **(a)** Serum-starved wild-type and RhoB KO cells were infected with ΔSopB *S.* Typhimurium and lysed with Laemmli buffer at 0.5, 1, 2, 3, 4 hpi. LC3, p62 and GAPDH were assessed by Western blotting. A representative experiment is shown. (**b**) Quantifications of relative ratios of LC3 and p62 to GAPDH as indicated. Data display means ± s.e.m. of assessed protein levels derived from three independent experiments, * p < 0.05, ** p < 0.01, two-way ANOVA.

**Supplementary Figure S5.**
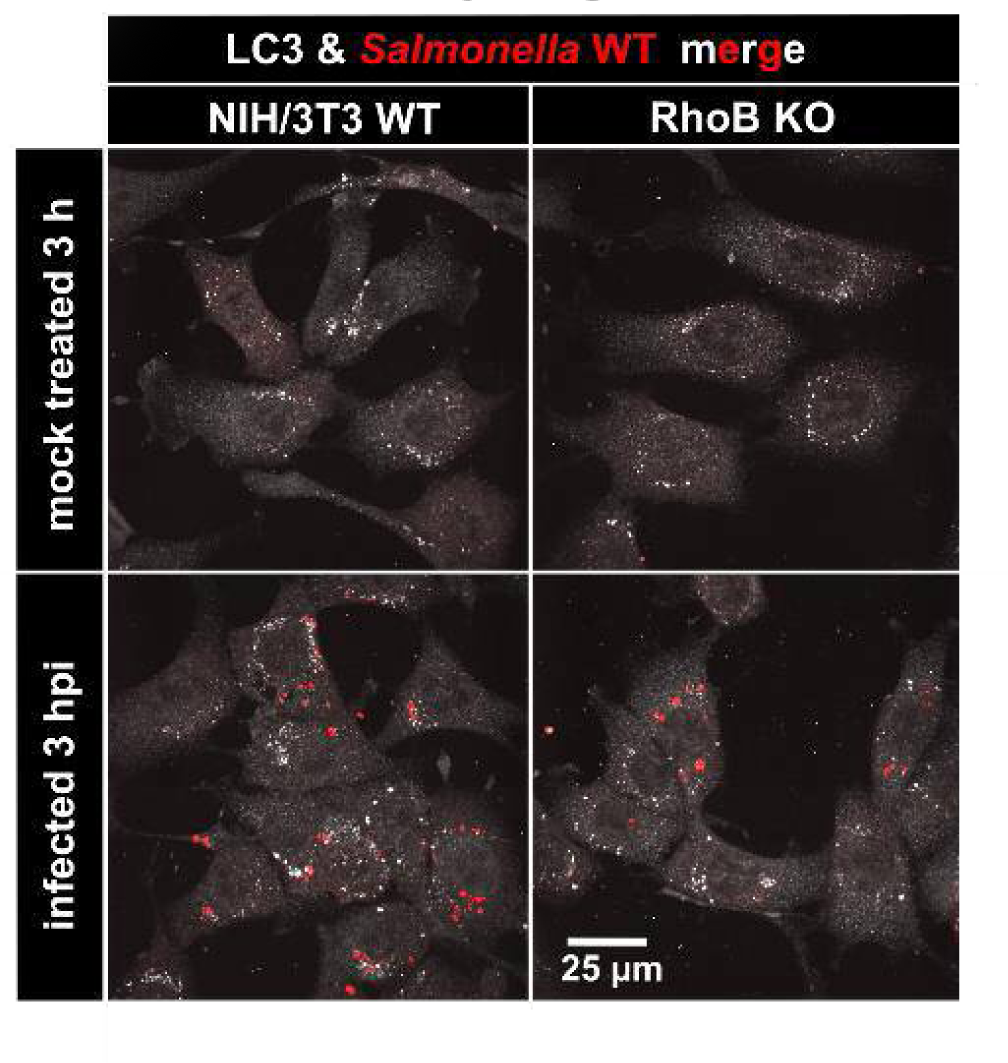
*Salmonella* invasion induces autophagosome formation in WT but not RhoB KO cells. NIH/3T3 wild-type and RhoB KO cells were mock-treated (top panel) or infected with *S.* Typhimurium wild-type expressing mCherry for 3 hours (bottom panel). Cells were fixed, immuno-stained for LC3 and acquired by spinning disk microscopy. Merged images show maximum intensity projections; LC3 is shown in white and *Salmonella* in red.

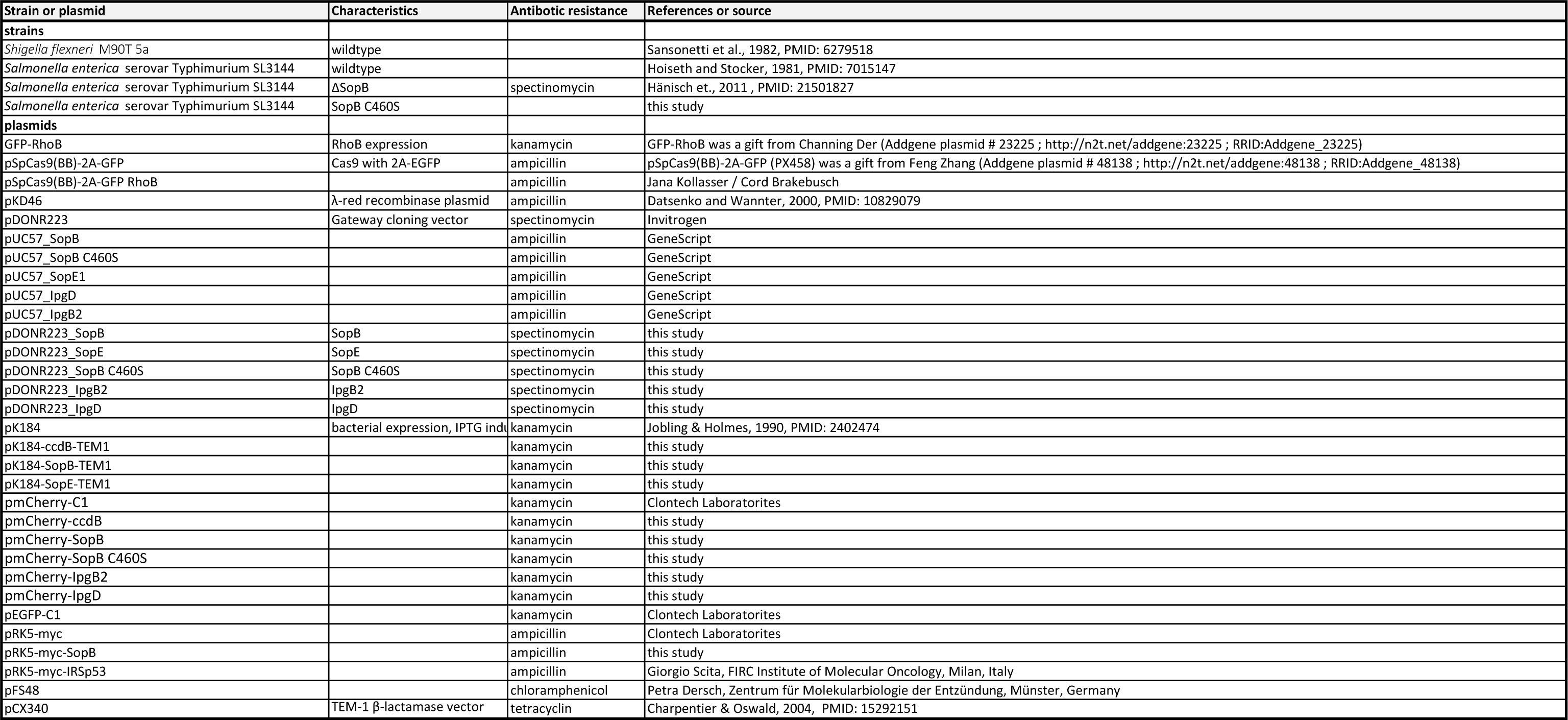

